# A genome-wide resource for the analysis of protein localisation in *Drosophila*

**DOI:** 10.1101/028308

**Authors:** Mihail Sarov, Chritiane Barz, Helena Jambor, Marco Y. Hein, Christopher Schmied, Dana Suchold, Bettina Stender, Stephan Janosch, Vinay K.J. Vikas, R.T. Krisnan, K. Aishwarya, Irene R.S. Ferreira, Radoslaw K. Ejsmont, Katja Finkl, Susanne Hasse, Philipp Kämpfer, Nicole Plewka, Elisabeth Vinis, Siegfried Schloissnig, Elisabeth Knust, Volker Hartenstein, Matthias Mann, Mani Ramaswami, K. VijayRaghavan, Pavel Tomancak, Frank Schnorrer

**Affiliations:** Max Planck Institute of Cell Biology and Genetics, Pfotenhauerstr. 108, 01307 Dresden, Germany; Muscle Dynamics Group, Max Planck Institute of Biochemistry, Am Klopferspitz 18, 82152 Martinsried, Germany; Department of Proteomics and Signal Transduction, Max Planck Institute of Biochemistry, Am Klopferspitz 18, 82152 Martinsried, Germany; Centre for Cellular and Molecular Platforms, National Centre for Biological Sciences, Tata Institute of Fundamental Research, Bald Bellary Road, Bangalore 560065, India; Heidelberg Institute of Theoretical Studies, Schloss-Wolfsbrunnenweg 35, 69118 Heidelberg, Germany; Department of Molecular Cell and Developmental Biology, University of California, Los Angeles, 610 Charles E. Young Drive, 5009 Terasaki Life Sciences Building, Los Angeles, CA 90095, USA; Institute of Neuroscience, Trinity College Dublin, Dublin 2, Ireland

## Abstract

The *Drosophila* genome contains >13,000 protein coding genes, the majority of which remain poorly investigated. Important reasons include the lack of antibodies or reporter constructs to visualise these proteins. Here we present a genome-wide fosmid library of ≈10,000 GFP-tagged clones, comprising tagged genes and most of their regulatory information. For 880 tagged proteins we have created transgenic lines and for a total of 207 lines we have assessed protein expression and localisation in ovaries, embryos, pupae or adults by stainings and live imaging approaches. Importantly, we can visualise many proteins at endogenous expression levels and find a large fraction of them localising to subcellular compartments. Using complementation tests we demonstrate that two-thirds of the tagged proteins are fully functional. Moreover, our clones also enable interaction proteomics from developing pupae and adult flies. Taken together, this resource will enable systematic analysis of protein expression and localisation in various cellular and developmental contexts.

## Impact statement

We provide a large-scale transgenic resource, which enables live imaging, subcellular localisation and interaction proteomics of selected gene products at all stages of *Drosophila* development.

## Introduction

With the complete sequencing of the *Drosophila* genome (Adams et al., 2000) genome-wide approaches have been increasingly complementing the traditional single gene, single mutant studies. This is exemplified by the generation of a genome-wide transgenic RNAi library (Dietzl et al., 2007) to systematically assess gene function in the fly or by the documentation of the entire developmental transcriptome during all stages of the fly’s life cycle by mRNA sequencing (Graveley et al., 2010). Furthermore, expression patterns were collected for many genes during *Drosophila* embryogenesis by systematic mRNA *in situ* hybrisation studies in different tissues (Hammonds et al., 2013; Tomancak et al., 2002; 2007). Particularly for transcription factors (TFs) these studies revealed complex and dynamic mRNA expression patterns in multiple primordia and organs during development (Hammonds et al., 2013), supposedly driven by specific, modular enhancer elements (Kvon et al., 2014). Furthermore, many mRNAs are not only dynamically expressed but also subcellularly localised during *Drosophila* oogenesis (Jambor et al., 2015) and early embryogenesis (Lécuyer et al., 2007). Together, these large-scale studies at the RNA level suggest that the activity of many genes is highly regulated in different tissues during development. Since the gene function is mediated by the encoded protein(s), the majority of proteins should display particular expression and subcellular localisation patterns that correlate with their function.

However, a lack of specific antibodies or live visualisation probes thus far hampered the systematic survey of protein expression and localisation patterns in various developmental and physiological contexts in *Drosophila.* Specific antibodies are only available for about 450 *Drosophila* proteins (Nagarkar-Jaiswal et al., 2015), and the versatile epitope-tagged UAS-based overexpression collection that recently became available (Bischof et al., 2013) is not suited to study protein distribution at endogenous expression levels. Collections of knock-in constructs are either limited to specific types of proteins (Dunst et al., 2015) or rely on inherently random genetic approaches, such as the large-scale protein-trap screens or the recently developed MiMIC (Minos Mediated Insertion Cassette) technology (Venken et al., 2011). The classical protein-trap screens are biased for highly expressed genes, and altogether recovered protein traps in 514 genes (Buszczak et al., 2007; Lowe et al., 2014; Morin et al., 2001; Quiñones-Coello et al., 2007). The very large scale MiMIC screen isolated insertions in the coding region of 1,862 genes, 200 of which have been converted into GFP-traps available to the community (Nagarkar-Jaiswal et al., 2015). Both approaches rely on transposons to mediate cassette insertion and require integration into an intron surrounded by coding exons for successful protein tagging. Thus, about 3,000 proteins, whose ORF is encoded within a single exon, cannot be tagged by these approaches. Together, this creates a significant bias towards trapping a particular subset of the more than 13,000 protein coding genes in the fly genome.

Hence, the *Drosophila* community would profit from a resource that enables truly systematic protein visualisation at all developmental stages for all protein coding genes, while preserving the endogenous expression pattern of the tagged protein. One strategy to generate a comprehensive resource of tagged proteins is to tag large genomic clones by recombineering approaches in bacteria and transfer the resulting tagged clones into animals as third copy reporter allele as was previously done in *C. elegans* (Sarov et al., 2012). In *Drosophila* it is possible to insert this tagged copy of the gene as a transgene at a defined position into the fly genome (Venken et al., 2006). The third copy reporter allele approach was used successfully with large genomic BAC or fosmid clones derived from the fly genome. It has been shown that these transgenes recapitulate the endogenous expression pattern of the gene in flies and thus likely provide a tagged functional copy of the gene (Ejsmont et al., 2009; Venken et al., 2009).

Here, we introduce a comprehensive genome-wide library of almost 10,000 C-terminally tagged proteins within genomic fosmid constructs. For 880 constructs, covering 826 different genes we generated transgenic lines, 765 of which had not been tagged by previous genetic trapping projects. Rescue experiments showed that two thirds of the tagged proteins are functional. We characterised the localisation pattern for more than 200 tagged proteins at various developmental stages from ovaries to adults by immunohistochemistry and by live imaging. This identified valuable markers for various tissues and subcellular compartments, many of which are detectable *in vivo* by live imaging. Together, this shows the wide range of possible applications and the potential impact this publically available resource will have on *Drosophila* research and beyond.

## Results

Our goal was to generate a comprehensive resource that allows the investigation of protein localisation and physical interactions for any fly protein of interest through a robust, generic tagging pipeline in bacteria, which is followed by a large-scale transgenesis approach (**Figure 1**). We based our strategy on a *Drosophila melanogaster* FlyFos library of genomic fosmid clones, with an average size of 36 kb, which covers most *Drosophila* genes (Ejsmont et al., 2009). Our two-step tagging strategy first inserts a generic ‘pre-tag’ at the C-terminus of the protein, which is then replaced by any tag of choice at the second tagging step, for example with a superfolder-GFP (sGFP) tag to generate the sGFP TransgeneOme clone library. These tagged clones are injected into fly embryos to generate transgenic fly-TransgeneOme (fTRG) lines, which can be used for multiple *in vivo* applications. (**Figure 1**).

**Figure 1.**
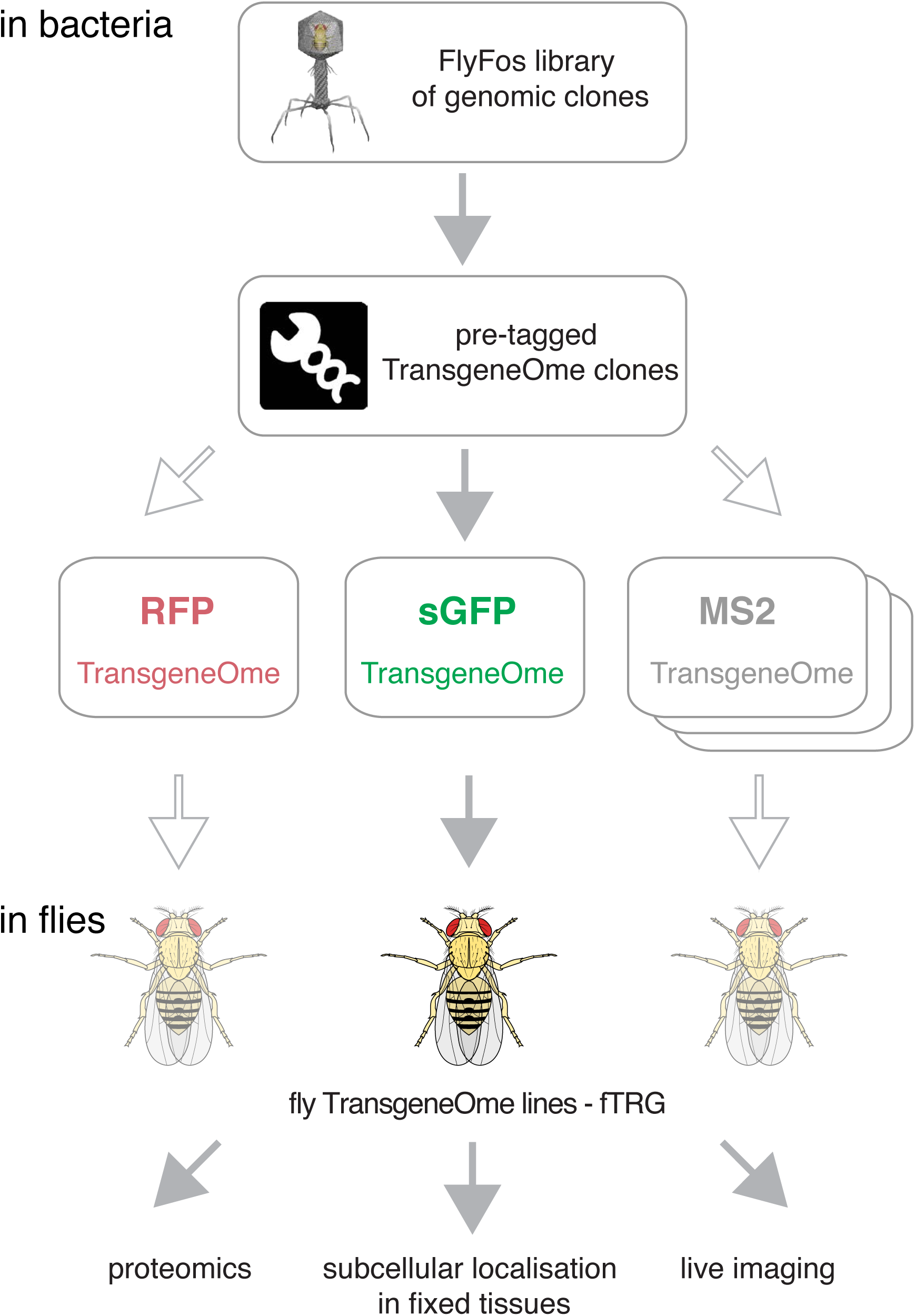
Overview of the tagging strategy and its applications. Liquid culture recombineering is used to insert a ‘pre-tagging’ cassette into FlyFos genomic clones in bacteria. This cassette can then be replaced by a simple, universal, recombineering reaction with any tag of choice, here, a superfolder GFP tag (sGFP) to generate the sGFP TransgeneOme clone library. These clones are transformed into flies generating transgenic FlyFos libraries that can be used for multiple *in vivo* applications. In the future, additional libraries with other tags can be generated easily.

## sGFP TransgeneOme – a genome-wide tagged FlyFos clone library

We aimed to tag all protein coding genes at the C-terminus of the protein, because a large number of regulatory elements reside within or overlap with the start of genes, including alternative promoters, enhancer elements, nucleosome positioning sequences, etc. These are more likely to be affected by a tag insertion directly after the start codon. Signal sequences would also be compromised by an inserted tag after the start codon. Additionally, the C-termini in the gene models are generally better supported by experimental data than the N-termini due to an historical bias for 3’ end sequencing of ESTs. Thus, C-terminal tagging is more likely to result in a functional tagged protein than N-terminal tagging, although we are aware of the fact that some proteins will be likely inactivated by addition of a tag to the C-terminus. Moreover, only about 1,400 protein coding genes contain alternative C-termini, resulting in all protein isoforms labelled by C-terminal tagging in almost 90 % of all protein coding genes.

In a series of pilot experiments we tested the functionality of several tagging cassettes with specific properties on a number of proteins (**Figure 2 Supplement 1, Table 1**). For the genome-wide resource we applied a two-step tagging strategy, whereby we first inserted a non-functional ‘pre-tagging’ cassette consisting of a simple bacterial selection marker, which is flanked with linker sequences present in all of our tagging cassettes. This strategy enables a very efficient replacement of the pre-tag by any tag of interest by homologous recombination mediated cassette exchange in bacteria. As fluorescent proteins and affinity tags with improved properties are continuously being developed, specific clones or the entire resource can be easily re-fitted to any new tagging cassette (**Figure 1**). For the genome-scale resource we selected a tagging cassette suitable for protein localisation and complex purification studies, consisting of the 2xTY1 tag as a flexible linker, the superfolder GFP coding sequence (Pédelacq et al., 2005), the V5 tag, followed by specific protease cleavage sites (for the PreScission- and TEV-proteases), the biotin ligase recognition peptide (BLRP) tag allowing for specific *in vivo* or *in vitro* biotinylation (Deal and Henikoff, 2010; Vernes, 2014), and the 3xFLAG tag (**Figure 2 Supplement 1**).

**Figure 2.**
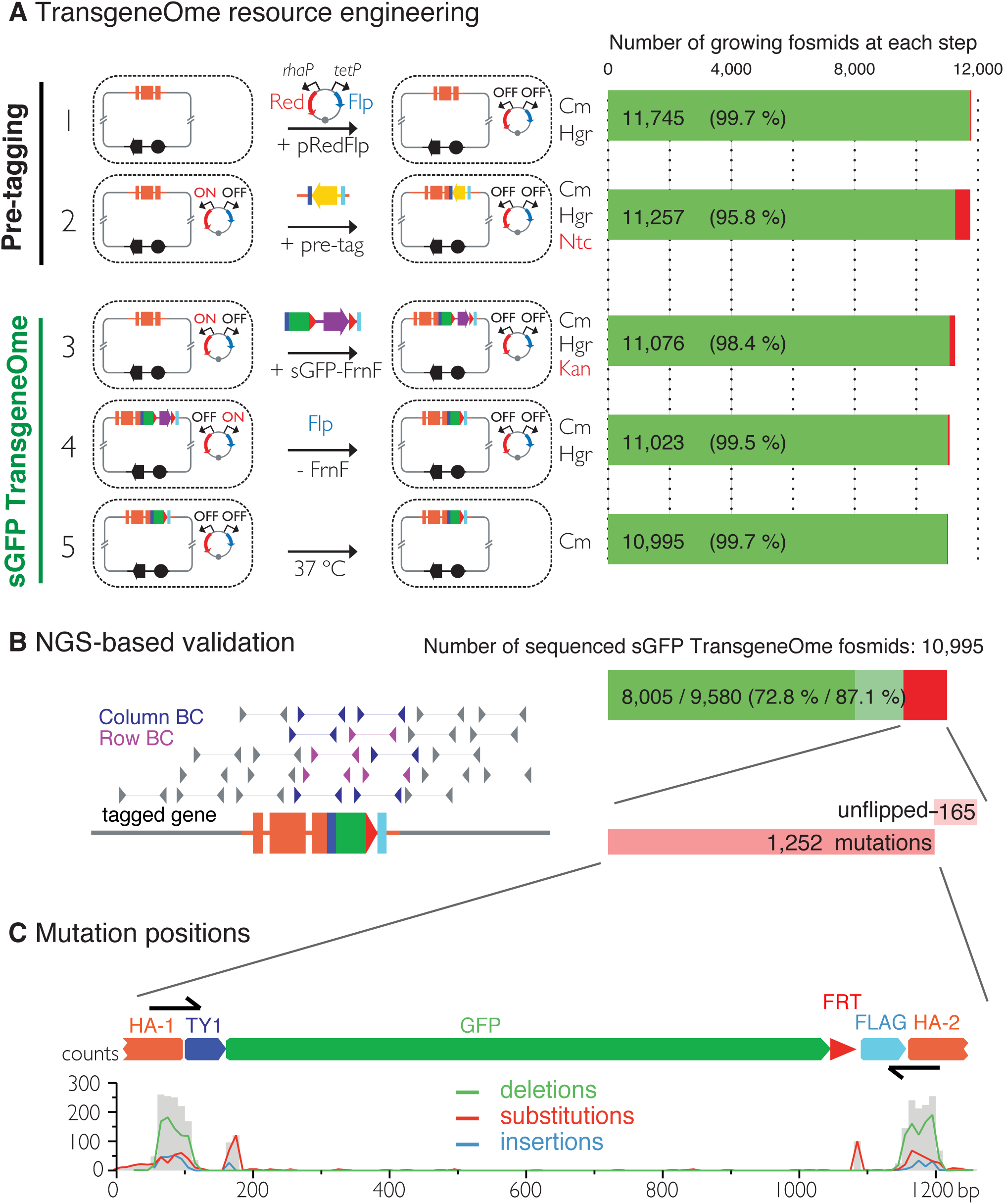
Generation of the TransgeneOme library. (**A**) TransgeneOme resource engineering. The steps of the recombineering pipeline are shown on the left with the success rate of each step indicated on the right (red colour denotes bacterial clones that did not grow). The *E. coli* cells are schematically represented with a dotted circle. With the first two steps the ‘pre-tagging’ cassette is inserted, which is replaced in the next three steps with the sGFP cassette to generate the sGFP TransgeneOme library. See text and methods section for details. (**B**) Next-generation-sequencing (NGS)-based validation of the sGFP TransgeneOme library. Schematic of the bar coding (BC) strategy of row and column pools is shown to the left and sequencing results to the right. Only the magenta and blue mate pairs contribute to the analysis of tag sequence. Fully sequenced tags within fosmids without point mutations are shown in solid green, clones without mutation in tagging cassette but incomplete coverage in light green and clones with mutation(s) or unflipped cassette are shown in red. (**C**) Statistics of the mutation distributions with deletions indicated by green, substitutions by red and insertions by blue lines. Note that most mutations reside within the primer sequences used as homology arms for recombineering (HA-1 and HA-2).

Of the 13,937 protein coding genes in the dmel5.43 genome assembly, 11,787 genes (84.6 %) were covered by a suitable fosmid from the original FlyFos library (Ejsmont et al., 2009), extending at least 2.5 kb upstream and 2.5 kb downstream of the annotated gene model. For picking clones, designing oligonucleotides for recombineering and for tracking all steps of the transgene engineering process, as well as for providing access to all construct sequences and validation data we used the previously developed TransgeneOme database (Sarov et al., 2012).

For high-throughput tagging of the *Drosophila* FlyFos clones, we developed an improved version or our previously applied high-throughput, 96-well format liquid culture recombineering pipeline (Ejsmont et al., 2011; Sarov et al., 2012). The high efficiency of recombineering in *E. coli* allowed for multi-step DNA engineering in 96-well format liquid cultures with single clone selection only at the last step. The specific pipeline consists of five steps (**Figure 2A**). First, the pRedFlp helper plasmid containing all genes required for homologous recombination as well as the Flippase-recombinase (under L-rhamnose and tetracyclin control, respectively) was introduced into *E. coli* by electroporation. Second, the ‘pre-tagging’ cassette containing only a bacterial antibiotic resistance gene was inserted by homologous recombination with gene-specific homology arms of 50 base pairs. Third, the sGFP-V5-BLRP tagging cassette, including an FRT-flanked selection and counter-selection cassette, was inserted to replace the ‘pre-tagging’ cassette. Since the linker sequences in the ‘pre-tagging’ cassette are identical to the tagging cassette, the tagging cassette was simply excised from a plasmid by restriction digest and no PCR amplification was required. This strongly reduced the risk of PCR induced mutations in the tagging cassette. Fourth, the selection marker was excised by the induction of Flippase expression. Fifth, the helper plasmid was removed by suppression of its temperature sensitive replication at 37 °C (Meacock and Cohen, 1980) and single clones were isolated from each well, by plating on selective solid agar plates.

All five steps of the engineering pipeline were highly efficient (between 95.8 and 99.7 %), resulting in an overall efficiency of 93.6 % or 10,995 growing cultures (**Figure 2A**). To validate the sequence of the engineered clones we developed a new next-generation-sequencing (NGS)-based approach (**Figure 2B**). In short, we pooled single clones from all 96-well plates into 8 rows and 12 columns pools and deep sequenced the barcoded and pooled mate pair libraries using the Illumina platform. The mate pair strategy allowed us to map the otherwise common tag coding sequence to a specific clone in the library and thus to verify the integrity of the tagging cassette insertion in the clones with single nucleotide resolution (see Material and Methods for details). When applied to the final sGFP TransgeneOme collection we detected no mutations for 9,580 constructs (87.1 %). 8,005 (72.8 %) of these clones had complete sequence coverage in the tag and thus represent the most reliable subset of the tagged library (**Figure 2B**). For 1,417 of the clones (12.8 %) one or more differences to the expected sequences were detected. The most common differences were point mutations, which cluster almost exclusively to the homology regions in the oligonucleotides used to insert the ‘pre-tagging’ cassette. This is suggestive of errors in the oligonucleotide synthesis but could also reflect polymorphisms in the genomic sequence of the homology arms. Another subset of point mutations clustered around the junctions between the homology arms and the rest of the tagging cassette, indicating an imprecise resolving of the homology exchange reaction in small subset of clones (**Figure 2C**). Finally, a small group of clones (165) still contained an unflipped selection cassette. The NGS results were confirmed by Sanger sequencing of the entire tag coding sequence for a subset of constructs (**Supplementary Table 1**).

Taken together, the sGFP TransgeneOme and our pilot tagging experiments resulted in 10,711 validated tagged clones, representing 9,993 different *Drosophila* genes. (**Table 1, Supplementary Table 1**). Moreover, the ‘pre-tagged’ TransgeneOme library is a versatile resource for generating fosmid clones with arbitrary tags at the C-terminus of the gene models.

**Table 1:**
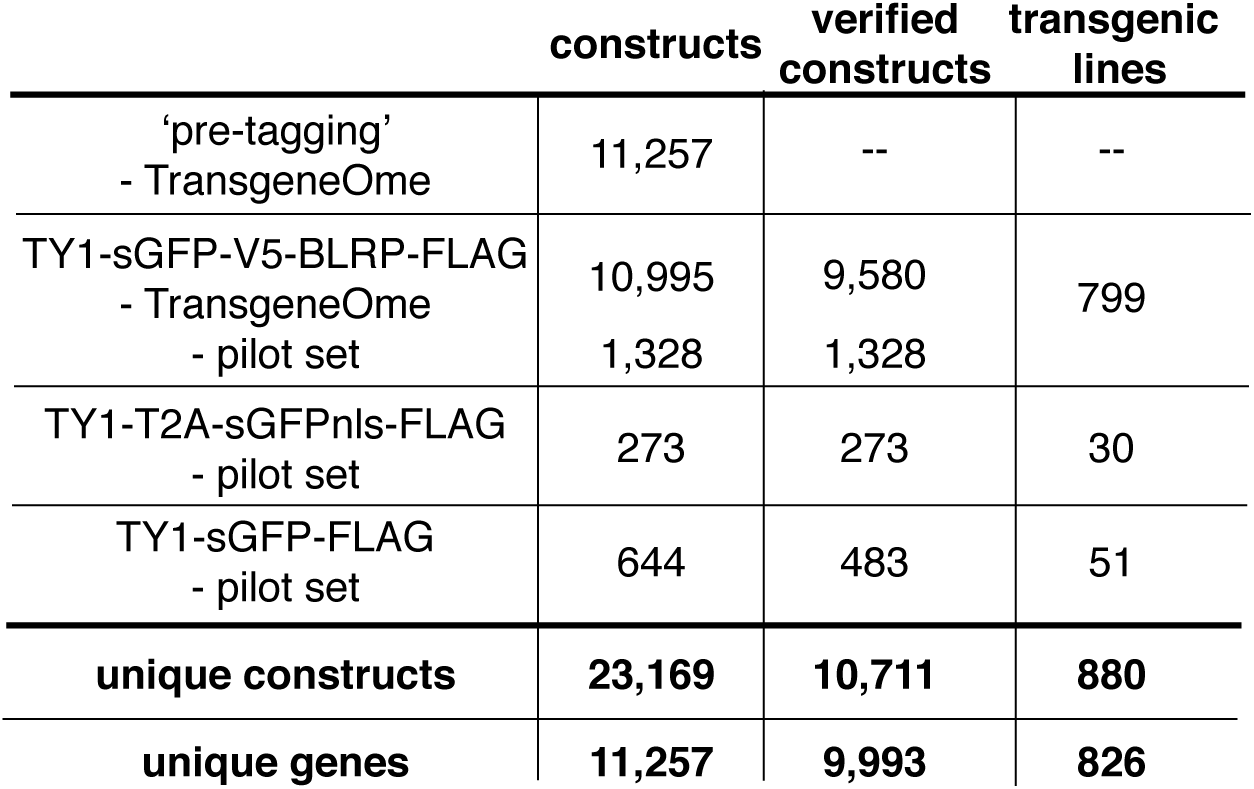
TransgeneOme constructs and fTRG lines - overview. Overview of TransgeneOme constructs generated and verified by sequencing for the different pilot sets and the genome-wide set, including the respective numbers of the transgenic fTRG lines generated.

Tagged constructs and transgenic lines
|  | constructs | verified<br>constructs | transgenic<br>lines |
| --- | --- | --- | --- |
| 'pre-tagging'<br>- TransgeneOme | 11,257 | -- | -- |
| TY1-sGFP-V5-BLRP-FLAG<br>- TransgeneOme | 10,995 | 9,580 | 799 |
| - pilot set | 1,328 | 1,328 |  |
| TY1-T2A-sGFPnls-FLAG<br>- pilot set | 273 | 273 | 30 |
| TY1-sGFP-FLAG<br>- pilot set | 644 | 483 | 51 |
| **unique constructs** | **23,169** | **10,711** | **880** |
| **unique genes** | **11,257** | **9,993** | **826** |

## Fly TransgeneOme (fTRG) – a collection of flies with tagged fosmids

We next established a pipeline to systematically transform the tagged TransgeneOme clones into flies. To efficiently generate fly transgenic lines we injected the tagged fosmid constructs into a recipient stock carrying the attP landing site VK00033 located at 65B on the third chromosome using a transgenic *nanos*-ΦC31 source (Venken et al., 2006). For some genes positioned on the third chromosome we injected into VK00002 located on the second chromosome at 28E to simplify genetic rescue experiments. In total, we have thus far generated lines for 880 tagged constructs representing 826 different genes (**Table 1**). These genes were partially chosen based on results of a public survey amongst the *Drosophila* community to identify genes for which there is the strongest demand for a tagged genomic transgenic line. 765 (87 %) of the newly tagged genes have not been covered by the previous protein-trap projects (**Supplementary Table 2**), hence, these should be particularly useful for the fly community. From our pilot tagging experiments, we made 51 lines for the 2xTY1-sGFP-3xFLAG tag and 30 lines for the 2xTY1-T2A-sGFPnls-FLAG transcriptional reporter. The majority of the lines (799) were generated with the versatile 2xTY1-sGFP-V5-Pre-TEV-BLRP-3xFLAG tag, used for the genome-wide resource (**Figure 2 Supplement 1, Table 1**). The collection of fly lines is called ‘tagged FlyFos TransgeneOme’ (fTRG) and all 880 fTRG lines have been deposited at the VDRC stock centre for ordering (http://stockcenter.vdrc.at).

To assess whether the tagged fosmids in our transgenic library are functional, we have chosen a set of 46 well-characterised genes, mutants of which result in strong developmental phenotypes. For most cases, we tested null or strong hypomorphic alleles for rescue of the respective phenotypes (embryonic lethality, female sterility, flightlessness etc.) with the tagged fosmid lines. More than two-thirds of the lines (31 of 46), including tagged lines of *babo, dlg1, dl, fat, Ilk, LanB1, numb, osk, rhea, sax, smo* and *yki* completely rescued the mutant phenotypes (**Figure 3**, **Supplementary Table 3**), demonstrating that the majority of the tagged fosmids are functional. Our rescue test set is biased towards important developmental regulators; 10 of the 15 genes that did not show a complete rescue are transcription factors with multiple essential roles during development, such as *esg, eya, odd, sna* and *salm.* Thus, their expression is likely regulated by complex cis-regulatory regions that may not be entirely covered by the available fosmid clone; for example wing-disc enhancers are located more than 80 kb away from the start of the *salm* gene (de Celis, 1999). Hence, we expect that a typical gene, which is embedded within many other genes in the middle of the fosmid clone, is more likely to be functional. Together, these data suggest that both the genome-wide tagged construct library and the transgenic fTRG library provide functional reagents that are able to substitute endogenous protein function.

**Figure 3:**
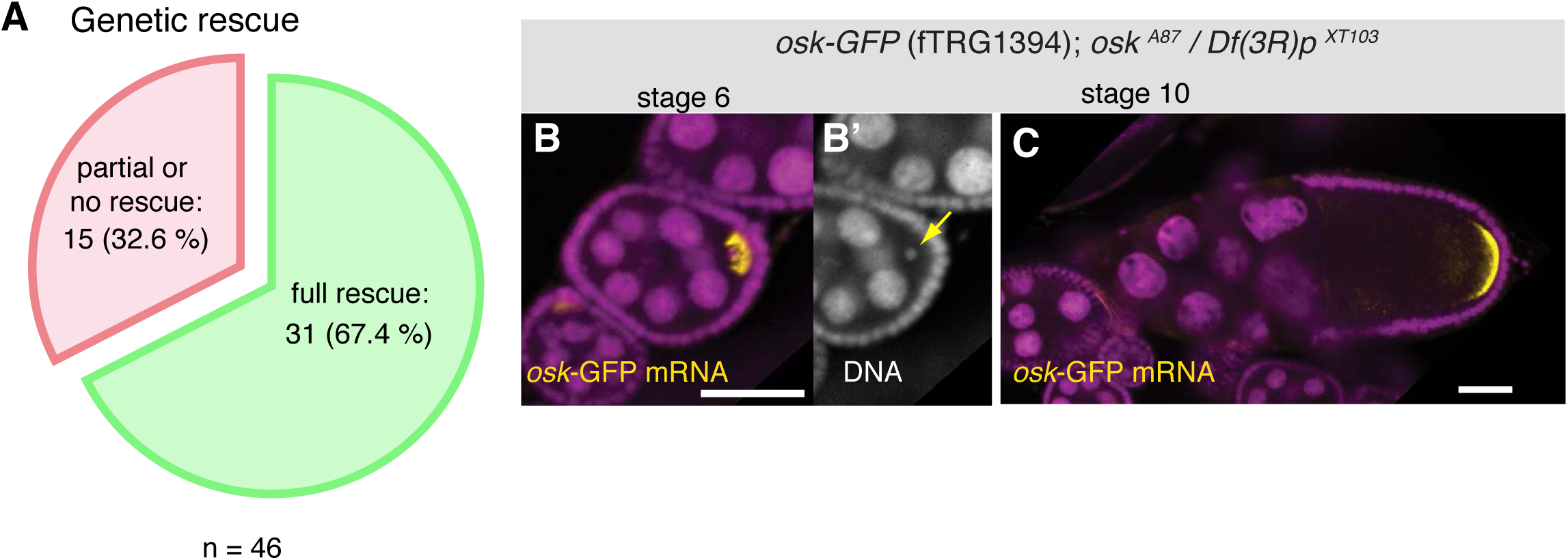
Functionality of the fTRG lines. (**A**) Genetic rescue statistics of null/strong mutant alleles for 46 selected fTRG lines. Note that more than two-thirds of the lines show a full rescue (see **Supplementary Table 3**). (**B**, **C**) *osk*-GFP mRNA (in yellow) expressed from fTRG1394 fully rescues egg-chamber development of an *osk* null allele (Jenny et al., 2006). *osk*-GFP mRNA enriches in the early oocyte (**B**, stage 6) and rescues the oogenesis arrest and the DNA condensation defect of the *osk* mutant (**B’**, yellow arrow). At stage 10 *osk*-GFP RNA enriches at the posterior pole (**C**) and produces sufficient protein to ensure proper embryogenesis. *osk*-GFP mRNA is shown in yellow, DAPI in magenta; scale bars indicate 30 μm.

## Expression of fTRG lines in the ovary

To demonstrate the broad application spectrum of our TransgeneOme library we analysed tagged protein expression and subcellular localisation in multiple tissues at various developmental stages. Germline expression in flies differs substantially from somatic expression, requiring particular basal promoters and often specialised 3’UTRs (Ni et al., 2011; Rørth, 1998). Therefore, we used ovaries to test the fTRG library and probed the expression of 115 randomly selected lines in germline cells versus somatic cells during oogenesis (**Figure 4A**). From the 115 lines 91 (79 %) showed detectable expression during oogenesis, with 45 lines being expressed in both, germ cells and the somatic epithelial cells (**Figure 4B, C** and **Supplementary Table 4**). 76 (66 %) fTRG lines showed interesting expression patterns restricted to subsets of cells or to a subcellular compartment (**Figure 4B - D**). For example, Tan-GFP is expressed in germline stem cells only, whereas the ECM protein Pericardin (Prc-GFP) is concentrated around the neighbouring cap cells and the transcription factor Delilah (Dei-GFP) is specifically localised to the nuclei of somatic stem cells, which will give rise to the epithelial cells surrounding each egg chamber (**Figure 4A, C**). In early egg chambers Reph-GFP is expressed in germ cells only, whereas the ECM protein Viking (Vkg-GFP) specifically surrounds all the somatic epithelial cells. Interestingly, the transcription factor Auracan (Ara-GFP) is only expressed in posterior follicle cells, whereas the putative retinal transporter CG5958 is only detectable in the squamous epithelial cells surrounding the nurse cells (**Figure 4C**).

**Figure 4:**
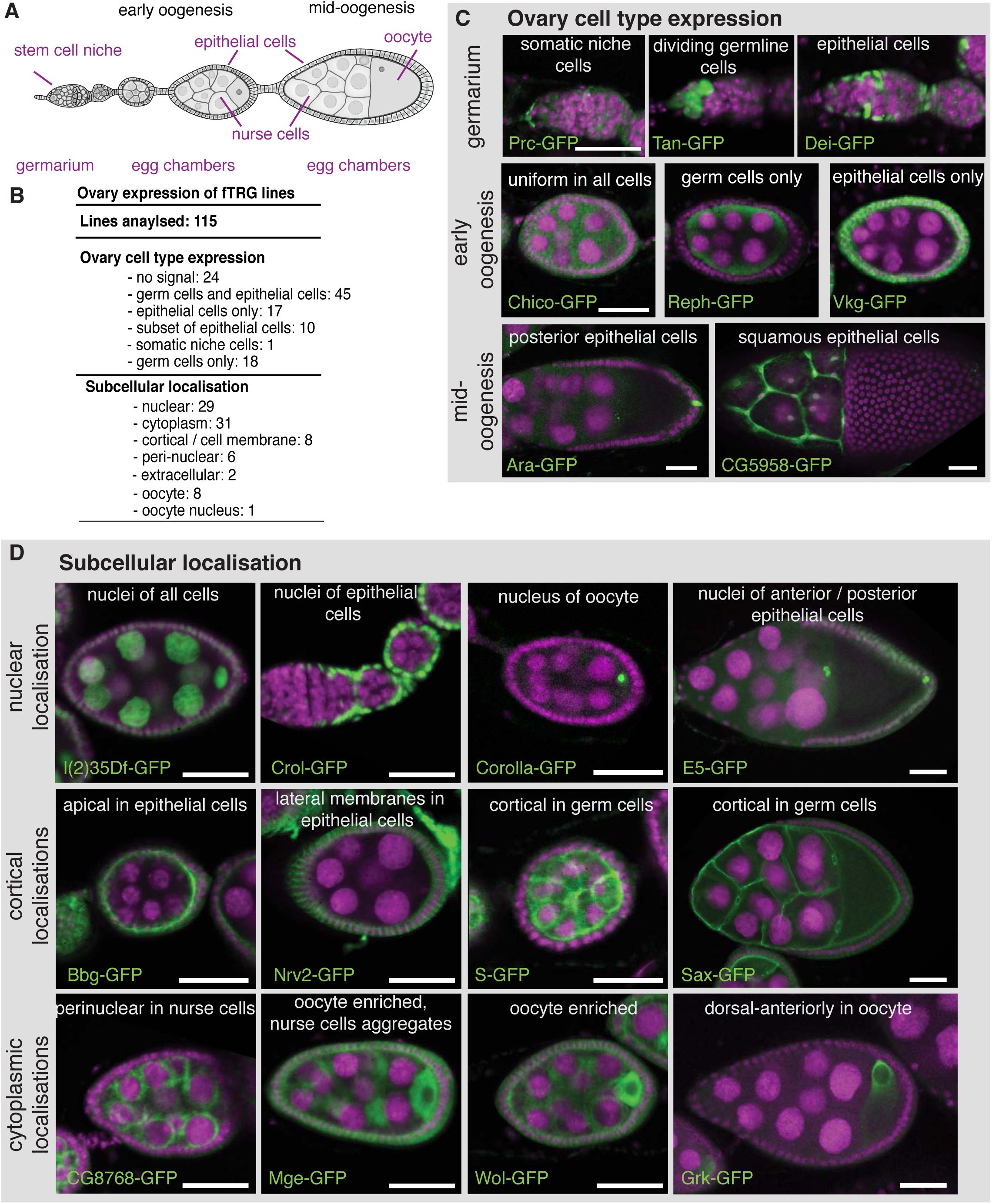
Expression of tagged transgenes in ovaries. (**A**) Schematic overview of oogenesis stages and cell types. (**B**) Summary of the identified expression patterns; see also **Supplementary Table 4**. (**C**) Selected examples for cell type specific FlyFos expression patterns at germarium, early- and mid-oogenesis stages visualised by anti-GFP antibody staining. (**D**) Selected examples of subcellular localisation patterns, highlighting nuclear, cortical and cytoplasmic patterns at different oogenesis stages. GFP is show in green, DAPI in magenta; scale bars indicate 30 μm.

We further investigated the subcellular localisation of the tagged proteins, which revealed a localisation for the RNA helicase l(2)35Df to all nuclei, whereas the predicted C_2_H_2_-Zn-finger transcription factor Crooked legs (Crol-GFP) is restricted to the nuclei of the epithelial cells (**Figure 4D**). Interestingly, Corolla-GFP is exclusively localised to the oocyte nucleus in early egg chambers. This is consistent with the function of Corolla at the synaptonemal complex attaching homologous chromosomes during early meiosis (Collins et al., 2014). In contrast, the uncharacterised homeobox transcription factor E5 (E5-GFP) is largely restricted to the nuclei of anterior and posterior epithelial cells (**Figure 4D**). Apart from nuclear patterns, we found a significant number of cortical localisations, including the well characterised Crumbs (Crb-GFP) (Bulgakova and Knust, 2009) and the PDZ-domain containing Big bang (Bbg-GFP) (Bonnay and Cohen-Berros, 2013) at the apical cortex of the epithelial cells, the Na^+^/K^+^ transporter subunit Nervana 2 (Nrv2-GFP) at the lateral epithelial membrane, and the EGF-signalling regulator Star (S-GFP) as well as the TGF-β receptor Saxophone (Sax-GFP) localised to the cortex or membrane of the germ cells **(Figure 4D** and **Supplementary Table 4**). Furthermore, we find a perinuclear enrichment for the uncharacterised predicted NAD binding protein CG8768, and oocyte enrichments for the Tom22 homolog Maggie (Mge-GFP) (Vaskova et al., 2000), glycosyltransferase Wollknäuel (Wol-GFP) (Haecker et al., 2008) and the TGF-α homolog Gurken (Grk-GFP), the latter with its well established concentration around the oocyte nucleus (Neuman-Silberberg and Schüpbach, 1993) (**Figure 4D**).

To test if genes expressed from FlyFos system also undergo normal post-transcriptional regulation we analysed the *osk-GFP* line, which was recently used as a label for germ granules (Trcek et al., 2015). *osk* mRNA is transcribed from early stages of oogenesis onwards in the nurse cell nuclei and specifically transported to the oocyte, where it localises to the posterior pole (Ephrussi et al., 1991; Kim-Ha et al., 1991). Only after the mRNA is localised, it is translated from stage 9 onwards (Kim-Ha et al., 1995). Indeed, fosmid derived *osk-GFP* mRNA localizes normally during all stages of oogenesis and its translation is repressed during mRNA transport, as Osk-GFP can only be detected at the posterior pole from stage 9 onwards (**Figure 4 Supplement 1A, B**). osk-GFP also fully rescues all aspects of an *osk* null allele (**Figure 3B, C**). Additionally, we discovered a post-transcriptional regulation for *corolla. corolla-GFP* mRNA is localised to the oocyte at stage 6 and Corolla-GFP protein is transported into the oocyte nucleus. However, despite the presence of the *corolla-GFP* mRNA at stage 8, Corolla protein is undetectable, suggesting either a translational block of the RNA or targeted degradation of the protein (**Figure 4 Supplement 1C - F**). Taken together, these expression and protein localisation data recapitulate known patterns accurately and identify various unknown protein localisations in various cell types during oogenesis, and thus emphasise the value of the fly TransgeneOme resource.

## *Live in toto* imaging of fTRG lines during embryogenesis

For many genes, the expression patterns at the mRNA level are particularly well characterised during *Drosophila* embryogenesis (Hammonds et al., 2013; Tomancak et al., 2002; 2007). However, *in situ* hybridisation techniques on fixed tissues do not visualise dynamics of expression over time and thus do not allow tracking of the expressing cells during development. As our tagging approach enables live imaging at endogenous expression levels we set out to test if *in toto* imaging using the SPIM (Selective Plane Illumination Microscopy) technology (Huisken et al., 2004) can be applied to the fly TransgeneOme lines. We pre-screened a small subset of lines (**Supplementary Table 5**) and selected the Na+/K+ transporter subunit Nrv2, as it shows high expression levels, for long-term time-lapse live imaging with a multi-view dual-side SPIM (Huisken and Stainier, 2007). During embryogenesis Nrv2 expression was reported in neurons (Sun et al., 1999) and glial cells (Younossi-Hartenstein et al., 2002). Interestingly, we find that Nrv2-GFP is already expressed from stage 11 onwards in most likely all cell types, where it localises to the plasma membrane, similar to the localisation in ovaries (**Figure 4D**). The expression level increases during stage 15 in all cells, with a particularly strong increase in the developing central nervous system (CNS) labelling the CNS and motor neuron membranes (**Figure 5, Supplementary Movie 1**^1^). These live *in toto* expression data are consistent with expression data of a recently isolated GFP trap in *nrv2* (Lowe et al., 2014), thus validating our methodology.

**Figure 5:**
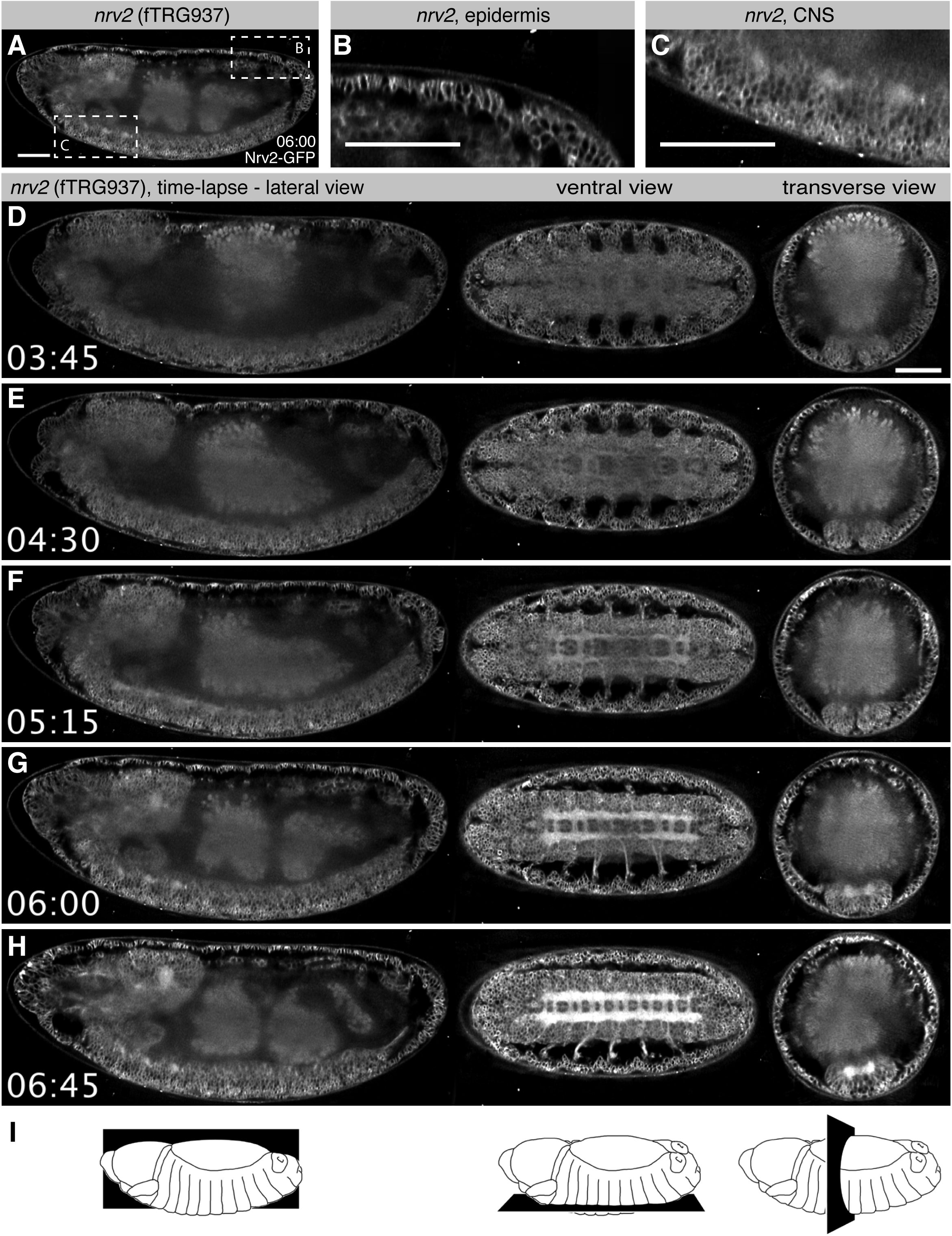
Live *in toto* imaging during embryogenesis with SPIM. (**A** - **C**) Nrv2-GFP protein is enriched in cell membrane of the epidermis and the CNS of late stage 16 embryos, as shown by a lateral section (**A**) and high magnifications of the posterior epidermis (**B**) and the ventral CNS (**C**). (**D** - **H**) Still image from a Nrv2-GFP time-lapse movie with lateral section views on the left, ventral sections in the middle and transverse sections on the right. Note that Nrv2-GFP is first expressed in the developing epidermal epithelial cells (**D**, **E**) and then becomes enriched in the CNS (**F** - **H**, see **Supplementary Movie 1**). (**I**) Schemes of the lateral, ventral and transverse section views through the embryo. Scale bars indicate 50 μm.

We wanted to extend our approach beyond highly expressed structural genes towards transcription factors that enable to follow cell lineages in the embryo. For this purpose we crossed the fTRG line of the homeobox transcription factor *gooseberry* (Gsb-GFP) to H2A-mRFPruby, which labels all nuclei, and recorded a two-colour multi-view dual-side SPIM movie. We find that Gsb-GFP is expressed in the presumptive neuroectoderm of the head region, labelling segmentally reiterated stripe-like domains at stage 10 (**Figure 5 Supplement 1A, B, Supplementary Movie 2**) as was described from fixed images (Gutjahr et al., 1993). Focusing on the deuterocerebral domain, we could reconstruct the delamination of two neuroblasts, which up-regulate Gsb-GFP while initiating their asymmetric divisions (**Figure 5 Supplement 1C - F**). It was possible to individually follow their neural progeny. Gsb-GFP expression also allowed us to directly follow the gradual down-regulation of Gsb-GFP in ectodermal cells that remained at the head surface after neuroblast delamination. As opposed to the neuroblasts, these cells, which give rise to epidermis, did not divide at all, or underwent only one further division (**Figure 5 Supplement 1G - J, Supplementary Movie 2**).

*gsb* is in part required for *gooseberry-neuro* (*gsb-n;* also called *gsb-d*) expression (He and Noll, 2013), which marks a defined set of brain neuroblasts in the embryo, with three *gsb-n* positive neuroblasts in the deuterocerebral domain (Urbach and Technau, 2003). We assume that these cells correspond to the three Gsb-GFP expressing neuroblasts visible in our live 4D stack. Notably expression in the Gsb-n-GFP fTRG line becomes detectable only at the end of germ-band extension (end of stage 11) in the developing CNS, where it lasts until stage 17 (**Figure 5 Supplement 2**, **Supplementary Movie 3**). Again this is consistent with published immuno-histochemistry data (Gutjahr et al., 1993; He and Noll, 2013). We conclude that our fly TransgeneOme library can be used for live *in toto* imaging, even for transcription factors expressed at endogenous levels. This will be of significant importance for ongoing efforts linking the transcription factor expression patterns of embryonic neuroblasts to the morphologically defined lineages that structure the larval and adult *Drosophila* brain (Hartenstein et al., 2015; Lovick et al., 2013; Pereanu, 2006).

## Expression of fTRG lines in the adult thorax

Cells in the embryo are generally rather small and thus not ideally suited to document subcellular protein localisation patterns. Thus, we decided to apply our TransgeneOme library to the adult thorax that contains some very large cells. We scored expression in the large indirect flight muscles (IFMs), in leg muscles, in visceral muscles surrounding the gut, in the gut epithelium, the tendon epithelium, the trachea and the ventral nerve cord including the motor neurons. In total, we found detectable expression in at least one tissue for 101 of 121 (83.5 %) analysed fTRG lines, thus creating a large number of valuable markers for cell types and subcellular structures (**Supplementary Tables 6, 7**).

The large IFMs are fibrillar muscles, which have a distinct transcriptional program resulting in their distinct morphology (Schönbauer et al., 2011). This is recapitulated by the expression of Act88F-GFP, which localises to the thin filaments of IFMs only (**Figure 6A - C**), whereas Mlp84B-GFP is not in IFMs but at the peripheral Z-discs of leg and visceral muscles only (**Figure 6D - F**), similar to the published localisation in larval muscle (Clark et al., 2007). We find various dotty patterns indicating localisation to intracellular vesicles; a particularly prominent example is Tango1-GFP in the midgut epithelium (**Figure 6G, H, Supplementary Table 6**). Tango1 regulates protein secretion in S2 cells, where it localises to the Golgi apparatus upon over-expression (Bard et al., 2006), suggesting that the pattern described here is correct. We find Par6 with the expected apical localisation in the proventriculus epithelium and in trachea (**Figure 6I, J**), whereas we identified a surprising pattern for the TRP channel Painless (Tracey et al., 2003). Pain-GFP is not only highly expressed in motor neurons (**Figure 6K**) but also in particular cells in the gut epithelium and most surprisingly, in the tendon cells to which the IFMs attach (**Figure 6L, M**). At this point we can only speculate that Pain might be involved in mechanical stretch-sensing in these cells. We have also tagged various ECM components, with LamininB1 (LanB1-GFP), LamininA (LanA-GFP) and BM40-SPARC resulting in the most prominent expression patterns. All three ensheath most adult tissues, particularly the muscles (**Figure 6N, O**, **Figure 6 Supplement 1A, B, E, F**). Interestingly, LanA-GFP and LanB1-GFP also surround the fine tracheal branches that penetrate into the IFMs, whereas BM40-SPARC is only detected around the large tracheal stalk and the motor neurons (**Figure 6P**, **Figure 6 Supplement 1C, D, G, H**). Finally, we also detected prominent neuromuscular junction (NMJ) markers; the IκB homolog Cactus shows a distinct pattern on leg muscles, visceral muscles and IFMs, the latter we could confirm by co-staining with the neuronal marker Futsch (**Figure 6 Supplement 1I - M**). Interestingly, such a NMJ pattern for Cactus and its binding partner Dorsal has been shown in larval body muscle by antibody stainings (Bolatto et al., 2003). Together, these results suggest that our fly TransgeneOme library provides a rich resource for tissue-specific markers in the adult fly that can routinely be used to visualise subcellular compartments in various tissues.

**Figure 6:**
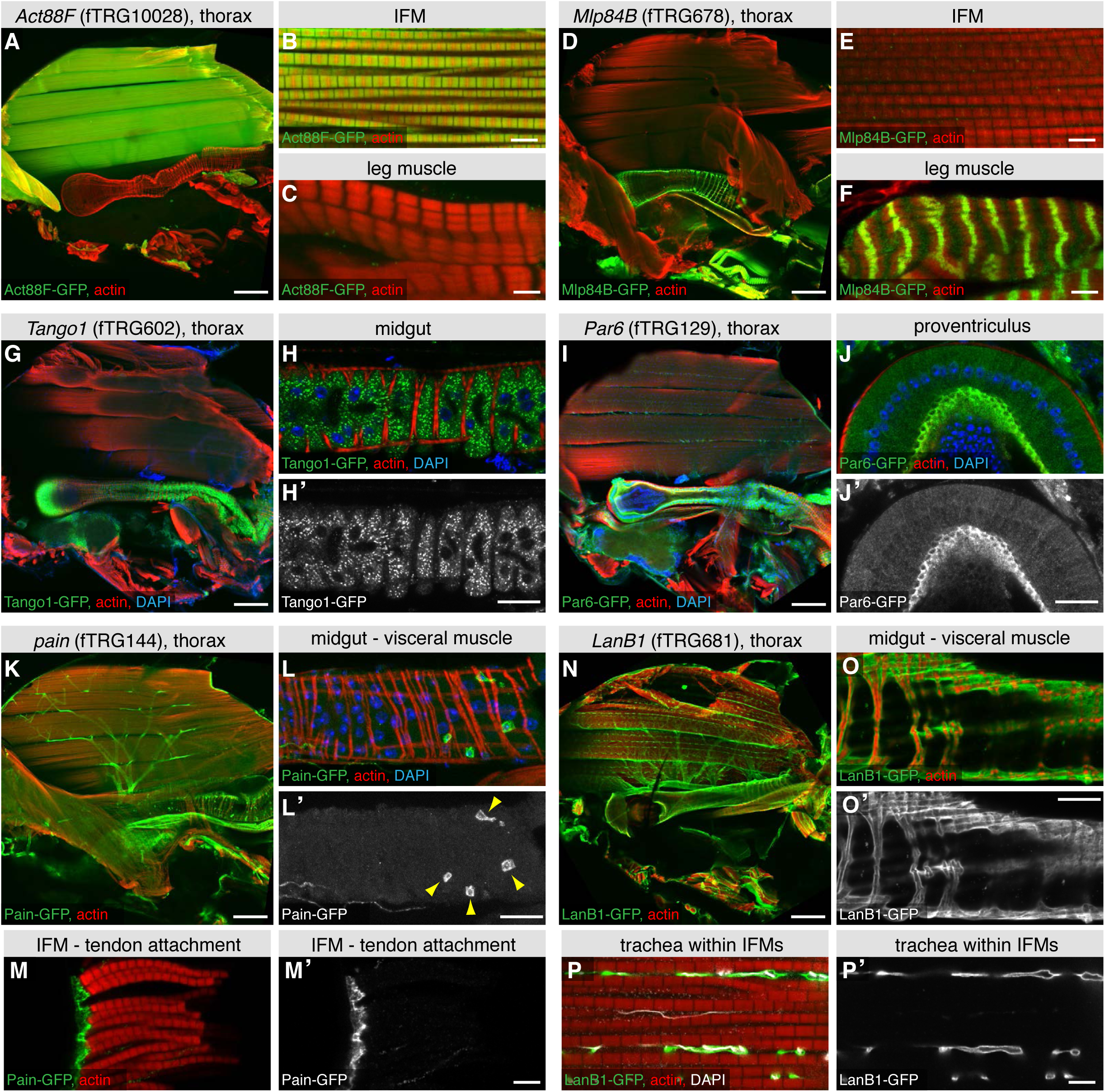
FlyFos expression in tissues of the adult thorax. Antibody stainings of the adult thorax with anti-GFP antibody (green) and phalloidin (red). (**A** - **F**) Act88-GFP expression is specific to the IFMs, where it labels the thin filaments (**B**), whereas Mlp84B specifically labels the Z-discs of leg muscles (**F**). (**G** - **J**) Tango1-GFP concentrates in a vesicle-like pattern in the gut epithelium (**H, H’**), whereas Par6-GFP is highly expressed in trachea (**I**) and the gut epithelium, where it concentrates at the apical membrane, as shown for a cross-section of the proventriculus (**J, J’**), nuclei are labelled with DAPI (blue). (**K** - **M**) Pain-GFP expression in the flight muscle motor neurons (**K**), cells close to the visceral muscles cells (possibly hemocytes) (**L, L’**) and tendon cells (**M, M’**). (**N** - **P**) LanB1-GFP labels the extracellular matrix surrounding the IFMs, the motor neurons and the trachea (**N**), as well as the visceral muscles (**O**). Even the finest trachea marked by UV auto-fluorescence (white) (**P**) are surrounded by LanB1-GFP (**P’**). Scale bars indicate 100 μm (A, D, I, K, N), 20 μm (H, J, L, O) and 5 μm (B, C, E, F, M, P).

To further validate the advantages of our TransgeneOme lines to label subcellular structures we imaged the large IFMs of the same 121 lines at high resolution. We found various markers for the thick filaments (e.g. the myosin associated protein Flightin, Fln-GFP) (Vigoreaux et al., 1993), for the myofibrils (e.g. the protein kinase Fray-GFP), the M-lines (e.g. the titin related protein Unc-89/Obscurin-GFP) (Katzemich et al., 2012), the Z-discs (e.g. CG31772-GFP) and the muscle attachment sites (e.g. Integrin-linked-kinase, Ilk-GFP). Furthermore, we identified markers for the T-tubules (e.g. Dlg1-GFP), for different vesicular compartments (e.g. the TGFβ receptor Baboon-GFP) and for mitochondria (CG12118) within the IFMs (**Figure 7, Supplementary Tables 6, 7**). Additionally, we documented the nuclear localisation in IFMs and leg muscles for a variety of fTRG proteins, including the uncharacterised homeodomain protein CG11617 and the C_2_H_2_ Zinc-fingers CG12391 and CG17912 (**Figure 7 Supplement 1A - C, E - G**); both of the latter result in flightless animals when knocked-down by muscle-specific RNAi (Schnorrer et al., 2010) suggesting that these genes play an essential role for IFM morphogenesis or function. Interestingly, the well characterised C_2_H_2_ Zinc-finger protein Hunchback (Hb) is only localised to leg muscle nuclei, but absent from IFMs suggesting a leg muscle-specific function of Hb (**Figure 7 Supplement 1G, H**).

**Figure 7:**
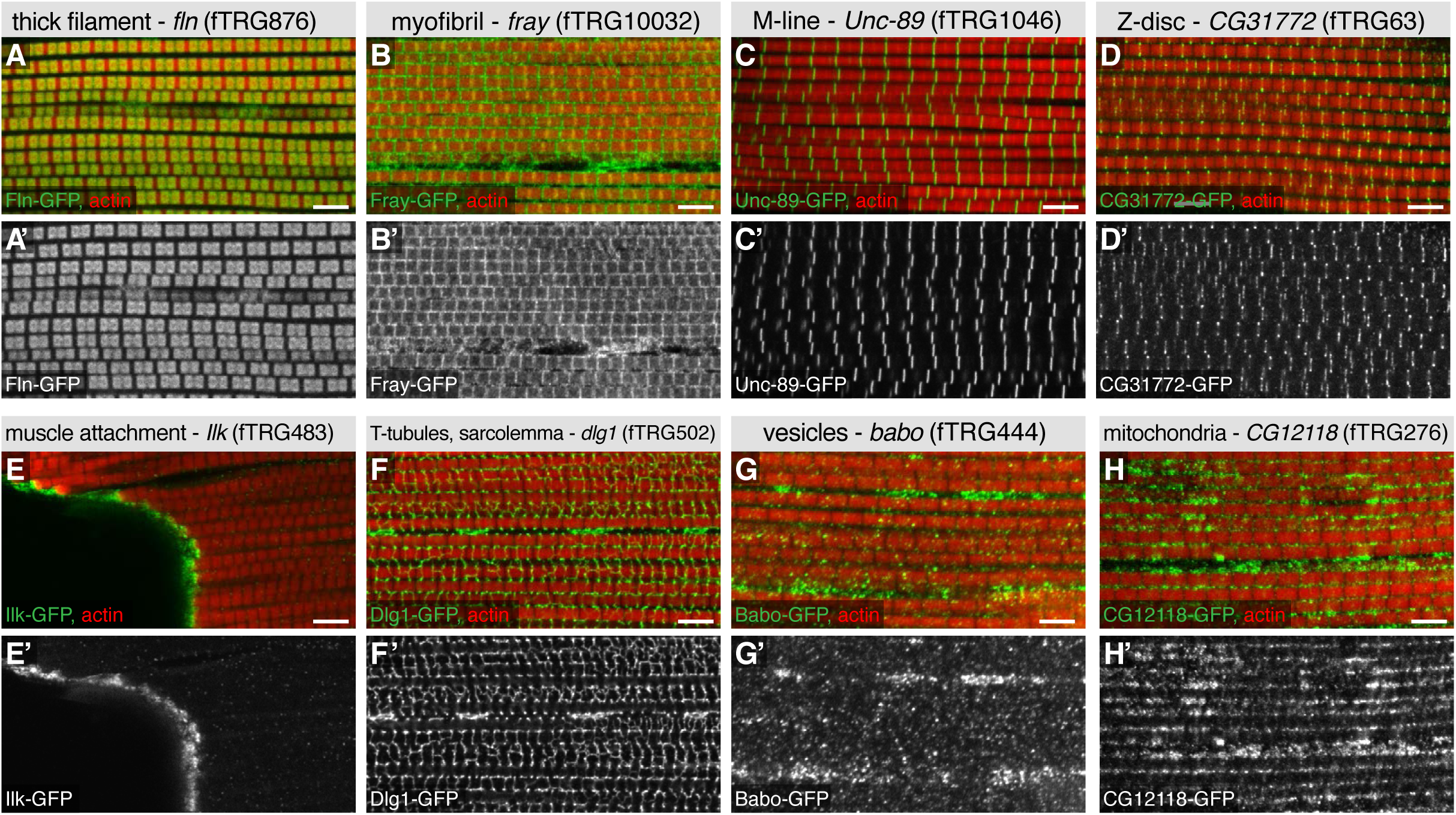
Subcellular patterns in adult flight muscles. Antibody stainings of the adult thorax with anti-GFP antibody (green or white in the single colour images) and phalloidin (red). (**A** - **D**) Localisation to specific myofibrillar sub-regions; Fln-GFP marks the thick filaments (**A**, **A’**), Fray-GFP surrounds the myofibrils with an enrichment at M-lines and Z-discs (**B**, **B’**), Unc-89-GFP marks only M-lines (**C**, **C’**) and CG31772-GFP only Z-discs (**D**, **D’**). (**E** - **H**) Ilk-GFP strongly concentrates at the muscle-tendon attachment sites (**E**, **E’**), Dlg1-GFP labels the T-tubular membranes (**F**, **F’**), Bab-GFP shows a dotty, vesicular pattern (**G, G’**) and CG12118 displays a mitochondrial pattern (**H**, **H’**). Scale bars indicate 5 μm.

However, differences between muscle types are not only controlled transcriptionally but also by alternative splicing (Oas et al., 2014; Spletter and Schnorrer, 2014; Spletter et al., 2015). To investigate if our tagging approach can be used to generate isoform-specific lines, we have chosen two prominent muscle genes, *mhc* and *rhea* (the fly Talin), both of which have predicted isoforms with different C-termini (**Figure 7 Supplement 2A, H**). Interestingly, we found that Mhc-isoforms K, L, M are expressed in IFMs and all leg muscles, however the predicted Mhc-isoforms A, F, G, B, S, V with the distal STOP codon are selectively expressed in visceral muscle and in a subset of leg muscles, however absent in adult IFMs (**Figure 7 Supplement 2B - G**). Even more surprisingly, while the long ‘conventional’ *rhea* (Talin) isoforms B, E, F, G show the expected localisation to muscle attachment sites in IFMs and leg muscles (Weitkunat et al., 2014), the short Talin-isoforms C and D do not localise to muscle attachment sites, but are selectively concentrated at costamers of leg muscles (**Figure 7 Supplement 2I - N**). Hence, our TransgeneOme library is ideally suited to label subcellular compartments and protein complexes, and in some cases can even distinguish between closely related protein isoforms.

## Expression of the fTRG lines in the living pupal thorax

An attractive application of the fly TransgeneOme library is live *in vivo* imaging. In the past, we had established live imaging of developing flight muscles in the pupal thorax using overexpressed marker proteins (Weitkunat et al., 2014). Here, we wanted to test, if live imaging of proteins at endogenous expression levels is also possible in the thick pupal thorax. We selected six fTRG lines for well established genes and indeed could detect expression and subcellular localisation for all of them using a spinning-disc confocal microscope either at the level of the pupal epidermis or below the epidermis, in the developing flight muscles, or both (**Figure 8**). The adducin-like Hts-GFP labels the cytoplasm of fusing myoblasts from 10 to 20 h APF (after puparium formation) and the developing SOPs (sensory organ precursors) with a particular prominent concentration in developing neurons and their axons (**Figure 8B - E**). In contrast, Dlg1-GFP localises to cell-cell-junctions of the pupal epidermis and to a network of internal membranes in the developing IFMs (**Figure 8F - I**) that may resemble developing T-Tubules, for which Dlg1 is a well established marker (Razzaq et al., 2001). Interestingly, the long isoforms of Talin-GFP (*rhea* isoforms B, E, F, G) are largely in the cytoplasm and at the cortex of the epidermal cells, with a marked enrichment in the developing SOPs at 10 to 20 h APF (**Figure 8J, K**). Further, Talin-GFP is strongly concentrated at muscle attachment sites of developing IFMs from 24 h onwards (**Figure 8L, M**) consistent with antibody stainings of IFMs (Weitkunat et al., 2014).

**Figure 8:**
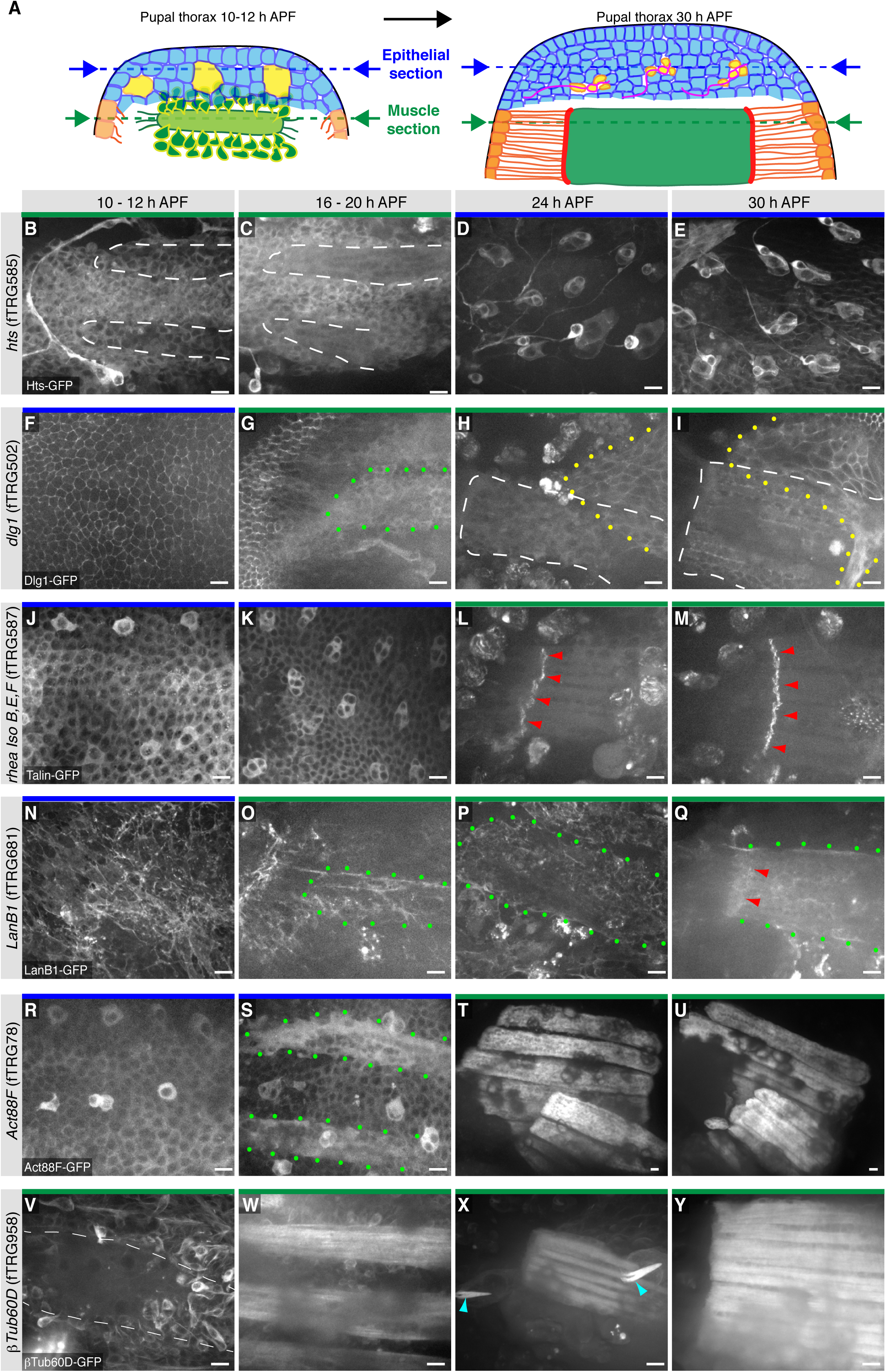
Live imaging of fTRG expression in living pupal thorax. (**A**) Schematic drawing of a 10 - 12 h (left) and a 30h pupal thorax (right). The developing epidermis is shown in blue, with the SOP precursors in yellow (developing neurons in red), the differentiating tendons are shown in orange, the myoblasts and muscle fibers in green, and the muscle-tendon junction in red. The schematic positions of the optical sections through epithelium and muscles are indicated with blue and green dotted lines, respectively. (**B** - **Y**) Live imaging of pupal thoraces at the indicated stages acquired with a spinning disc confocal (except **S** and **T**, which were acquired with a two-photon microscope). Blue bars above the image indicate epithelial sections and green bars indicate muscle sections (as explained in **A**). Hts-GFP is expressed in fusing myoblasts (**B**, **C**) and strongly in developing SOPs (**D**, **E**). Dlg1-GFP labels the epithelial junctions (**F**), internal muscle structures (green dots, **G**) and an unidentified additional developing epithelium (yellow dots, **H**, **I**). Talin-GFP is higher expressed in developing SOPs (**J**, **K**) and strongly localised to the muscle-tendon junction from 24h APF (red arrowheads, **L**, **M**). LanB1-GFP localises to the basal side of the developing epithelium (**N**) and surrounds the forming muscle fibers (green dots, **O** - **Q**) with a slight concentration at the muscle-tendon junction at 30h APF (red arrowheads, **Q**). Act88F-GFP weakly labels the developing epithelium, with a slight concentration in the SOPs until 20h APF (**R**, **S**) and very strongly marks the IFMs from 24h onwards (**T**, **U**). βTub60D-GFPis expressed in the fusing myoblasts (**V**, **W**) and also labels the microtubule bundles in the developing muscle fibers (**X**, **Y**) and hair cells of the developing sensory organs (light blue arrow heads in **X**). Scale bars indicate 10 μm.

The dynamics of the extracellular matrix is little described thus far as very few live markers existed. Hence, we tested our LamininB1 fosmid and found that LanB1-GFP is readily detectable within the developing basement-membrane basal to the epidermal cells of the pupal thorax at 10h APF (**Figure 8N**). It also labels the assembling basement-membrane around the developing IFMs from 16 to 30h APF without a particularly obvious concentration at the muscle attachment sites (**Figure 8O - Q**). To specifically visualise the developing IFMs we chose Actin88F, which is specifically expressed in IFMs and a few leg muscles (Nongthomba et al., 2001). We find that the Act88F-GFP fTRG line indeed very strongly labels the IFMs from about 18 h APF but is also expressed in the developing pupal epidermis again with an enrichment in the forming SOPs from 10 to 20 h APF (**Figure 8R - U**). The latter is not surprising as Act88F-lacZ reporter has been shown to also label the developing wing epithelium (Nongthomba et al., 2001), again suggesting that our fTRG line recapitulates the endogenous expression pattern. We finally tested the βTub60D fTRG-line, as βTub60D was reported to label the myoblasts and developing myotubes in embryonic and adult muscles (Fernandes et al., 2005; Leiss et al., 1988; Schnorrer et al., 2007). Indeed, we detect βTub60D-GFP in fusing myoblasts and the developing IFMs, with particularly prominent label of the microtubule bundles at 24h APF (**Figure 8V - Y**). In addition, βTub60D-GFP also strongly marks the developing hairs of the sensory organs of the pupal epidermis (**Figure 8X**, see also **Supplementary Movie 6**).

In order to test, if the fly TransgeneOme lines and the sGFP-tag are indeed suited for long-term live imaging we chose Act88F-GFP and βTub60D-GFP and imaged the developing IFMs for more than 19 h with a two-photon microscope using an established protocol for over-expressed markers (Weitkunat and Schnorrer, 2014). For both proteins we can detect strongly increasing expression after 18h APF in the developing IFMs, with Act88F-GFP being restricted to the myotubes and the developing myofibrillar bundles (**Supplementary Movie 4**, **Figure 8 Supplement 1A - F**) whereas βTub60D-GFP also labels the fusing myoblasts and is largely incorporated into prominent microtubule bundles (**Supplementary Movie 6**, **Figure 8 Supplement 1L - Q**).

As photo bleaching was no serious problem in these long movies we also recorded movies at higher time and spatial resolution. We labelled the developing IFMs with Act88F-GFP and the myoblasts with a *him*-GAL4, UAS-palm-Cherry and acquired a 3D stack every two minutes using a spinning disc-confocal. This enabled us to visualise single myoblast fusion events in developing IFMs of an intact pupa (**Supplementary Movie 5**, **Figure 8 Supplement 1G - K**). The six dorsal longitudinally oriented IFMs develop from three larval template muscles to which myoblasts fuse to induce their splitting into six myotubes (Fernandes et al., 1991). Using high resolution imaging of βTub60D-GFP we find that most myoblasts fuse in the middle of the developing myotube during myotube splitting, with prominent microtubules bundles located at the peripheral cortex of the splitting myotube (**Supplementary Movie 7, Figure 8 Supplement 1R**). These prominent microtubules bundles are then relocated throughout the entire developing myotube (**Supplementary Movie 7**, **Figure 8 Supplement 1S - V**). Taken together, these live imaging data suggest that many of the fTRG lines will be well suited for high resolution live imaging of dynamic subcellular protein localisation patterns in developing *Drosophila* organs. This will strongly expand the set of live markers available for research in flies.

## FlyFos library as bait for proteomics

For the proper composition, localisation and *in vivo* function of most protein complexes the expression levels of the individual components are critical (Rørth et al., 1998; Tseng and Hariharan, 2002). Hence, the TransgeneOme library would be an ideal experimental set-up to purify protein complexes from different developmental stages using endogenous expression levels of the bait protein. In principle, all the small affinity tags (TY1, V5, FLAG) (**Figure 2 Supplement 1**) can be used for complex purifications. The presence of precision and TEV cleavage sites even allow two-step purifications. For proof of principle experiments, we selected four tagged proteins as baits: Ilk, Dlg1, Talin and LanB1, and analysed two different developmental stages. In each case we homogenised hundred 24 to 48 h pupae and hundred adult flies per experiment and mixed the cleared lysate with a GFP antibody matrix to perform single step affinity enrichment and mass-spec analysis modified from the QUBIC protocol (Hein et al., 2015; Hubner et al., 2010; Keilhauer et al., 2014). Each affinity-enrichment was performed in triplicate and intensity profiles of all identified proteins were quantified in a label-free format by running all 30 purifications consecutively on the same Orbitrap mass-spectrometer and analysing the data with the MaxQuant software suite (Cox and Mann, 2008; Cox et al., 2014) (**Supplementary Table 8**). Interestingly, enriching Ilk-GFP from both developing pupae and adult flies recovered the entire Ilk, PINCH, Parvin, RSU-1 complex (**Figure 9**), which had previously been purified *in vitro* from *Drosophila* S2 cells (Kadrmas et al., 2004) and mammalian cells (Dougherty et al., 2005; Tu et al., 2001) giving us confidence in our methodology. We also successfully enriched Talin-GFP from pupae or adults, however did not identify an obvious strong and specific binding partner (**Figure 9**, **Supplementary Table 8**). In contrast, we identified Mesh as a novel interactor of Dlg1 from pupae and adult flies. Mesh colocalises with Dlg1 at septate junctions of the embryonic *Drosophila* midgut, however a molecular interaction of both proteins was not established (Izumi et al., 2012). Finally, we purified the laminin complex by pulling on LanB1, which recovered LanB2 and LanA roughly stoichiometrically, both from pupae and adult flies, as had been found in cell culture experiments (Fessler et al., 1987), showing that extracellular matrix complexes can also be purified from *in vivo* samples with our methodology. In summary, these data demonstrate that interaction proteomics with the fly TransgeneOme library can confirm known interaction partners and discover novel *in vivo* complex members, making the system attractive for a variety of biochemical applications.

**Figure 9:**
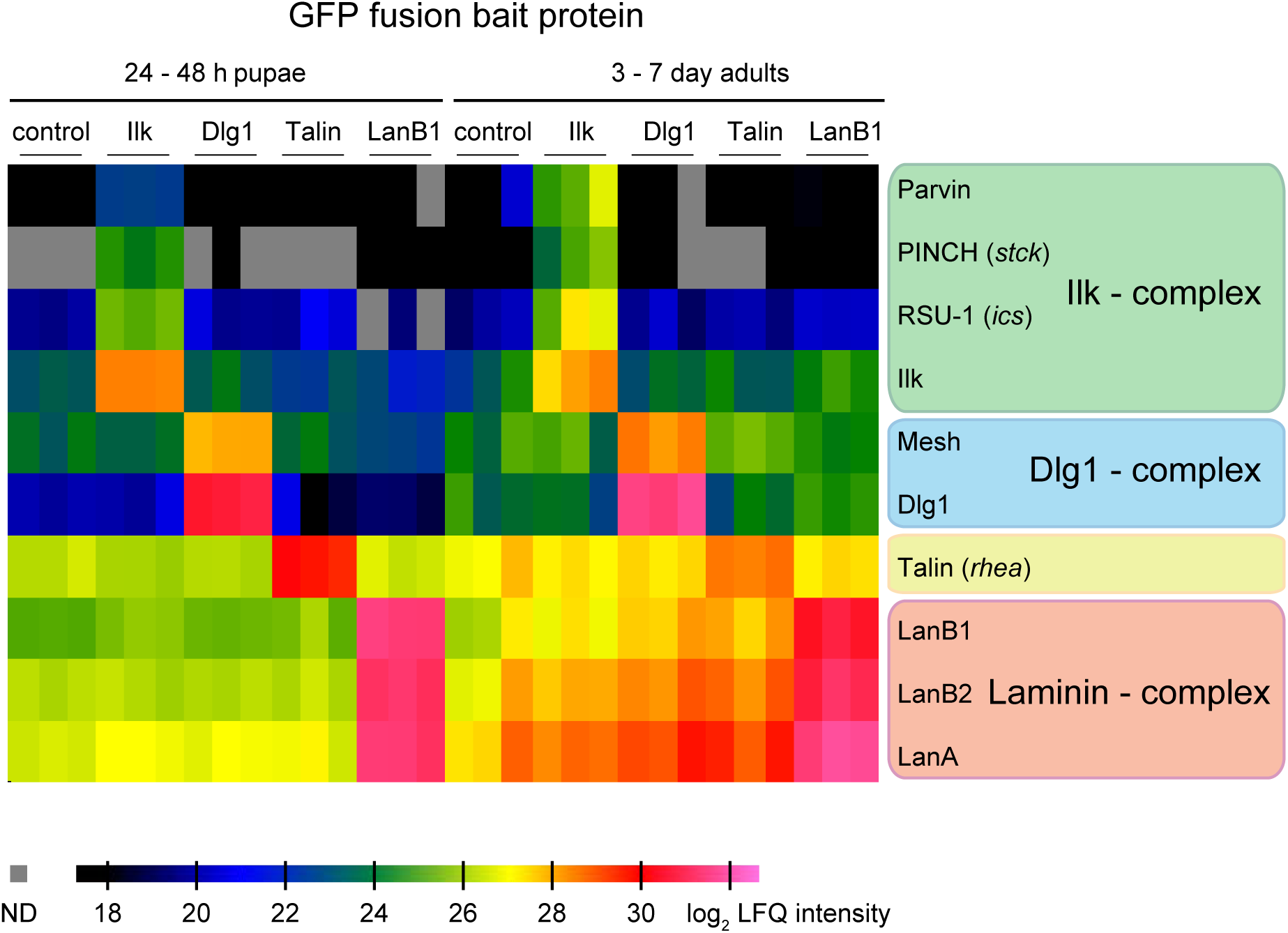
Proteomics with fTRG bait proteins. GFP pull-downs from Ilk-GFP, Dlg1-GFP, Talin-GFP, LanB1-GFP and control pupal (left) or adult fly (right) protein extracts were analysed by mass spectrometry and MaxQuant. The label-free quantification (LFQ) intensities are colour coded. They allow the comparison of protein amounts across samples and correlate with absolute protein amounts. Baits and specific interactors are characterized by quantitative enrichment over the negative controls and samples with non-interacting baits.

## Discussion

The TransgeneOme resource presented here adds new powerful component into the arsenal of tools available to the *Drosophila* research community. It complements the genetic resources for gene disruption and localisation (Buszczak et al., 2007; Lowe et al., 2014; Morin et al., 2001; Nagarkar-Jaiswal et al., 2015; Quiñones-Coello et al., 2007; St Johnston, 2012; Venken and Bellen, 2012) with a comprehensive genomescale library that does not suffer the biases of random mutagenesis. Analogously to the powerful MiMIC system (Nagarkar-Jaiswal et al., 2015) the TransgeneOme resource is versatile and can be adapted to the developments in tag chemistry and to various specialised applications. Although the resource is designed to study behaviour of proteins, it can for example be easily converted into a toolkit for live imaging of mRNAs. By designing a tagging cassette with an array of MS2 binding sites (Forrest and Gavis, 2003) the existing ‘pre-tagged’ TransgeneOme can be converted into an MS2-tagged TransgeneOme by a simple liquid culture recombineering step in bacteria (**Figure 1**).

However, any new TransgeneOme has to be transformed into flies and this process still represents a significant bottleneck. We present here an optimised protocol for transgenesis of fosmid size clones into *Drosophila melanogaster.* It took three years and four dedicated technicians to generate the 880 fly lines presented in this study. Although, the systematic transgenesis is a continuing process in our laboratories, the value of the TransgeneOme collection is highlighted by the fact that any specific set of genomically tagged gene clones is now available. These can be efficiently transformed by in house transgenesis of *Drosophila* labs around the world using the optimised protocol presented here. For that purpose we will not only deposit the sGFP TransgeneOme but also the ‘pre-tagged’ TransgeneOme collection at Source Biosciences from which any subset of clones can be conveniently ordered (http://www.sourcebioscience.com/).

One caveat of designed expression reporters is the necessity to place the tag into a defined position within the gene model. We chose to generate our ‘pre-tagged’ collection at the most commonly used C-terminus predicted by the gene model, thus labelling most isoforms. In a few cases a tag at the C-terminus will inactivate the protein, however such a reagent can still be useful for visualising the protein. This has been demonstrated for a number of sarcomeric GFP-traps, some of which lead to lethality when homozygous, yet result in interesting localisation patterns when heterozygous (Buszczak et al., 2007; Morin et al., 2001 and F.S., unpublished observation). For particular genes, it will be useful to tag differential protein isoforms, which in some cases can be done by tagging alternative C-termini, as shown here for *Mhc* and *rhea.* However, tagging a particular isoform requires a very informed construct design, which cannot easily be automated at the genome-scale.

Genome engineering is experiencing a tremendous growth with the introduction of CRISPR/Cas technology and it will be only a matter of time before a larger collection of precisely engineered fusion proteins at endogenous loci will become available in flies. However to date, such examples are still limited to a few genes (Baena-Lopez et al., 2013; Gratz et al., 2014; Port et al., 2014; Zhang et al., 2014), which had been carefully picked and were individually manipulated with custom-designed, gene-specific tools. It remains to be tested which proportion of such engineered loci will be fully functional and thus potentially superior to the fTRG collection. Having a transgenic third allele copy, as is the case in our TransgeneOme collection, might even be advantageous, if the tagging interferes with protein function, because the TransgeneOme lines still retain two wild-type endogenous gene copies. In some cases, addition of GFP might destabilise the protein, regardless of Nor C-terminal fusion, as recently shown for the Engrailed protein (Sokolovski et al., 2015). However, our ability to detect the protein product in the vast majority of our tagged lines argues that this could be a relatively rare, gene specific phenomenon. Nevertheless, caution should be taken with respect to protein turnover dynamics of any tagged protein.

Together, the FlyFos library, the fly TransgeneOme library and the fTRG collection of strains, enable genome-scale examination of expression and localisation of proteins comparable with the high-throughput mRNA *in situ* screens (Tomancak et al. 2002, Tomancak et al. 2007). Our data for tagged Oscar protein show that fosmid reporters can in principle recapitulate all aspects of gene expression regulation at transcriptional and post-transcriptional levels. It will be particularly interesting to combine the spatial expression data of mRNAs with that of proteins. Since many transcripts show subcellular localisation in various developmental contexts (Jambor et al., 2015; Lécuyer et al., 2007), the question arises whether RNA localisation generally precedes localised protein activity. Systematic examination of protein patterns expressed from localised transcripts in systems such as the ovary will provide a genome-scale overview of the extent and functional role of translational control. At the tissue level, the patterns of mRNA expression may be different from the patterns of protein expression, for example due to translational repression in some cells or tissue specific regulation of protein stability, as shown here for the Corolla protein. The combined mRNA and protein expression patterns may therefore uncover a hidden complexity in overall gene activity regulation and the fTRG lines will help to reveal these combinatorial patterns in a systematic manner.

The fTRG lines faithfully recapitulate gene expression patterns in ovaries, embryos, larvae, pupae and adults suggesting that they can be used to visualise proteins in every tissue during the life cycle of the fly. This includes adult tissues such as the flight or leg muscles, which thus far had not been subjected to systematic protein expression and localisation studies. However, due to their size and the conservation of the contractile apparatus these tissues are particularly attractive to study with this new resource. In general, antibody or FISH stainings with a single standard anti-tag reagent are easier to optimise, compared to antibody stainings or mRNA *in situ* with gene specific antibodies/probes. This simplicity makes it possible to explore the expression of the available genes across multiple tissues, as has been done for the *rab* collection (Dunst et al., 2015). Such an approach is orthogonal to the collections of expression data generated thus far, in which many genes were examined systematically but only at particular stages or in certain tissues, i.e. embryos or ovaries (Jambor et al., 2015; Lécuyer et al., 2007; Tomancak et al., 2007). We are confident that the analysis of regularly studied as well as less explored *Drosophila* tissues will be stimulated by the fTRG collection.

When protein expression levels are sufficiently high, the fusion proteins can be visualised by live imaging approaches in intact animals. It is difficult to estimate the absolute expression levels required for live visualisation, as this depends on the imaging conditions, the accessibility and transparency of the tissue and importantly on the observed protein pattern. A strongly localised protein can result in a very bright local signal, such as Talin or Ilk at the muscle attachment sites, compared to a protein homogenously distributed throughout the entire cell. In particular for tissues, such as the adult legs, antennae or the adult fat body, which are difficult to dissect and stain without losing tissue integrity, these live markers should be enormously beneficial.

One important limitation for examining the pattern of protein expression is the accessibility of the tissue of interest for imaging. We have shown that light sheet microscopy can be used to image the dynamics of tagged protein expression throughout embryogenesis. We further demonstrated that two-photon microscopy can be applied to study protein dynamics during muscle morphogenesis in developing pupae. Other confocal or light sheet based imaging paradigms could be adapted for *in toto* imaging of living or fixed and cleared specimen from other life cycle stages. Establishing standardized protocols for preparation, staining and imaging of *Drosophila* stages, isolated tissues and organs will be necessary to realise the full potential of the fTRG collection.

Protein interaction data in fly are available from a number of studies (Formstecher et al., 2005; Giot et al., 2003; Guruharsha et al., 2011). These results were generated using yeast-two-hybrid, or overexpression in a tissue culture system, followed by affinity-purification and mass spectrometric analysis. Despite high-throughput, these approaches face the problem that the interacting proteins might not be present at the same place within a cell, or not even co-expressed in a developing organism. This is circumvented by affinity purifications of endogenously expressed proteins, which thus far at genome-scale was only reported from yeast (Gavin et al., 2002; Ho et al., 2002; Krogan et al., 2006). In higher organisms, BAC-based systems, which are closely related to our fosmid approach, elegantly solved these issues, as shown by a recent human interactome study (Hein et al., 2015).

The collection of transgenic flies covers currently only about 10 % of the available tagged fosmids. Expanding the collection to include most genes of the genome and importantly characterising the transgenes-encoded proteins by imaging in various biological contexts is best achieved by spreading the clones and transgenic lines amongst the community of researchers using *Drosophila* as a model system. Therefore, all transgenic lines are available from the VDRC stock collection. Despite the expanding CRISPR-based genome engineering technologies, the fTRG collection will continue to be an important resource for the fly community, in particular, if the full functionality of certain fTRG lines has been demonstrated, as we did here for a selection of important developmental regulators. As with many genome-scale resources it is typically easier to produce them than to fully characterise and exploit their potential. Comprehensive generation of thousands of transgenes and their thorough analysis takes time; it took us 4 years to assemble the collection presented here. Development of protocols and techniques to image these collections of tagged lines and assembling open access databases to share the data needs to continue and will eventually become useful also for the characterisation of resources whose production began only recently.

Wangler, Yamamoto and Bellen convincingly argued that the *Drosophila* system remains an indispensable model for translational research because many essential fly genes are homologs of Mendelian disease genes in humans (Wangler et al., 2015). Yet, even after decades of research on fruit flies only about 2000 of the estimated 5000 lethal mutations have been investigated. Resources like ours will therefore provide essential functional information about gene expression and localisation in *Drosophila* tissues that can serve as a starting point for the mechanistic understanding of human pathologies and their eventual cures.

## Materials and Methods

### TransgeneOme clone engineering

Fosmids were engineered as described previously (Ejsmont et al., 2009; 2011), except for the inclusion of the ‘pre-tagging’ step in the genome-wide TransgeneOme set. All tagging cassettes were generated from synthetic DNA and cloned into R6K carrying plasmids, which require the presence of the *pir* gene product for replication (Metcalf et al., 1996). The *pir* gene is not present in the FlyFos library host strain, thereby ensuring near-complete lack of background resistance in the absence of the correct homologous recombination event.

Details of the recombineering steps are as follows (**Figure 2A**): Step 1. The *E. coli* cells containing a FlyFos clone covering the gene locus of interest are transformed with the pRedFlp plasmid, containing the genes necessary for the homologous recombination and the Flp recombinase under independently inducible promoters. Step 2. Next, a ‘pre-tagging’ cassette carrying an antibiotic resistance gene (NatR, nourseothricin resistance) surrounded by regions of homology to all specific tagging cassettes (**Figure 2 Supplement 1**) and flanked by gene specific homology arms is electroporated as linear DNA fragment produced by PCR. By combination of induced (L-rhamnose) pRedFlp homologous recombination enzyme action and strong selection with a cocktail of three antibiotics (one to maintain the fosmid (chloramphenicol, Cm), one to maintain the pRedFlp (hygromycin, Hgr) and nourseothricin (Ntc) to select for the inserted fragment) the electroporated linear ‘pre-tagging’ fragment becomes inserted in front of the STOP codon of the gene of interest. Step 3. The ‘pre-tagging’ cassette is exchanged for a cassette of the chosen tag coding sequence including an FRT flanked selection / counter selection marker (rpsL-neo). This cassette is now universally targeting the homologous sequences shared by the tagging and pre-tagging cassettes and is produced in bulk by restriction enzyme mediated excision from a plasmid. Note that in this way, no PCR induced mutations can be introduced at this step. Step 4. Upon Flp induction (with anhydrotetracycline) the rpsL-neo cassette is excised, leaving a single FRT site, positioned in frame with the tag coding sequence. Step 5. Finally, the recombineering plasmid is removed from the cells containing the engineered fosmids by inhibition of its temperature sensitive origin of replication and release from Hgr selection. The cells are plated on a selective chloramphenicol agar plate from which a single colony is picked and further validated.

### NGS-based validation of the TransgeneOme clones

For NGS-based resource validation single colonies for each TransgeneOme clone were picked into 96-well plates and individual wells of all 96-well plates were pooled into 8 rows and 12 columns pools. Fosmid DNA was isolated from these pools, and mate pair fragment libraries were prepared and sequenced on an Illumina platform. First, adapters and low quality sequences were trimmed with Trimmomatic0.32. (parameters: ILLUMINACLIP:NexteraPE-PE.fa:2:30:10 LEADING:3 TRAILING:3 SLIDINGWINDOW:4:15 MINLEN:36). Second, in order to detect un-flipped fosmid sequences (where the FLP-mediated excision of selection cassette failed), the read pairs were mapped with Bowtie2 (Langmead and Salzberg, 2012) against the unflipped tag sequence and the genome. If any read of the mate of the pair mapped to the un-flipped sequence while the second mate mapped to the genome consistent with the estimated mate pair insert size of 3000 bp ± 1000 bp, the fosmid was flagged a unflipped and was not further analysed. Third, in order to identify mutations in the tag and in the immediate genomic surrounding (± 1000 bp), the NGS reads were mapped against the fosmid references that included the flipped tag. The Bowtie2 was set to report only hits where both reads of the pairs map concordantly to the insert size in the tag and in the genome (parameters: -I 2200 -X 3700 --rf --no-discordant --no-unal --no-mixed). This mapping was further filtered by deleting PCR duplicated read pairs with samtools1.1 *rmdup* (Li et al., 2009). Mutations were identified by utilising SNP calling implemented in FreeBayes (Garrison and Marth, 2012) using the standard filters and *vcffilter* to eliminate reported SNPs with scores < 20. Finally, in the last step, the information of the row and column pools were compared and summarized using a custom C-program that read the results of the SNP calling and the Bowtie mappings and counted the coverage for each read pair anchored in tag sequence with at least 20 bp. To correct for random PCR or sequencing errors the reported SNPs were compared for the row and column pools of each fosmid and SNPs occurring in both pools with coverage of 3 or more reads were considered as real.

### *Drosophila* stocks and genetic rescue experiments

Fly stocks were maintained using standard culture conditions. All crosses were grown at 25°C unless otherwise noted. Most of the fly mutant or deficiency strains for the rescue experiments were obtained from the Bloomington *Drosophila* Stock Center and if located on X- or 2^nd^ chromosome crossed together with the respective fTRG line. If the mutant gene was located on the 3^rd^ chromosome it was recombined with the fTRG line. Rescue was generally tested in trans-heterozygotes as indicated in **Supplemental Table 3**. The rescue for 6 genes *(bam, fat, mask, rap, RhoGEF2* and *yki*) was done by others, who communicated or published the results (**Supplementary Table 3**). For rescue of flightlessness a standard flight test was used (Schnorrer et al., 2010).

### Generation of transgenic FlyFos (fTRG) lines

Most TransgeneOme fosmid clones were injected into the y[1], w[*], P{nos-phiC31\int.NLS}X; PBac{y+-attP-3B}VK00033 (BL-32542). This stock has white eyes and no fluorescent eye markers, which would interfere with screening for the red fluorescent eye marker used in the FlyFos clones (Ejsmont et al., 2009). A few fosmid clones were also injected into y[1], w[*], P{nos-phiC31\int.NLS}X; PBac{y+-attP-3B}VK00002, with the attP site located on the 2^nd^ chromosome. The *osk-GFP* fosmid was injected into attP40.

#### Detailed injection protocol

A) Bacterial culture of fosmid clones: 1. Inoculate 2 ml LB-medium plus chloramphenicol (Cm 25 μg / ml) with fosmid clone and grow over night at 37 °C. 2. Dilute to 10 ml (9 ml LB-medium + Cm and 1 ml bacterial culture) and add 10 μl 10 % arabinose (final concentration 0.01 %) to induce the fosmid to high copy number. 3. Grow at 37 °C for 5 h and collect the pellet by 10 min centrifugation at 6000 rpm. Pellet can be stored at -20 °C.

B) Preparation of fosmid DNA: Use the HiPure Plasmid Miniprep Kit from Invitrogen (order number: K2100-03) according to the supplied protocol (MAN0003643) with following modifications: before starting: pre-warm the elution buffer (E4) to 50 °C; step 4: incubate the lysate for 4 min at room temperature; step 5: incubate 4 min on ice before centrifuging at 4 °C for 10 min; step 8: add 850 μl elution buffer (pre-warmed to 50 °C) to the column; step 9: add 595 μl isopropanol to the elution tube, centrifuge 20 min at 4 °C; wash pellet with 800 μl 70 % ethanol; centrifuge for 2 min; step 12: air dry the pellet for 4 min. Add 20 μl EB-buffer (Qiagen) to the pellet and leave at 4 °C overnight to dissolve without pipetting to avoid shearing of the DNA. Do not freeze the DNA. Adjust the concentration to 250 ng / μl and centrifuge 5 min at full speed before injections. Do not inject DNA older than one week.

C) Embryo injections: Collect young embryos (0 - 30 min) on an agar plate, bleach away the chorion, wash and collect the embryos on a cellulose filter (Whatman 10409814). Align the embryos, transfer them to a glued slide and dry them with silica gel for 10 - 15 min (Roth T199.2). Cover the embryos with Voltalef 10S oil (Lehmann & Voss) and inject the prepared fosmid DNA using a FemtoJet set-up (Eppendorf 5247). The injected DNA should be visible within the embryo. Incubate the injected embryos for 48 h at 18 °C in a wet chamber and collect the hatched larvae with a brush. Cross the surviving mosaic adults individually to *y, w* males or virgins.

### Immuno-stainings

Ovaries: sGFP-protein detection in egg-chambers was done as previously described (Dunst et al., 2015). Detection of the *oskar-GFP* mRNA was performed with a *gfp-*antisense probe (Jambor et al., 2014) and co-staining of *osk* mRNA and Osk protein was done as previously described (Jambor et al., 2011) using a *gfp*-antisense probe and a rabbit anti-GFP antibody (1:1000, ThermoFisher).

Adult thoraces: Antibody stainings of adult thoraces, including flight, leg and visceral muscles, were done essentially as described for adult IFMs (Weitkunat and Schnorrer, 2014). Briefly, thoraces from young adult males were fixed for 15 min in relaxing solution (20 mM phosphate buffer, pH 7.0; 5 mM MgCl_2_; 5 mM EGTA, 5 mM ATP, 4 % PFA) + 0.5 % Triton X-100, cut sagittally with a sharp microtome blade and blocked for 1 hour at room temperature with 3 % normal goat serum in PBS-0.5% Triton X-100. Samples were stained with primary antibodies overnight at 4 °C (rabbit anti-GFP 1:2000 Amsbio; mouse anti-Futsch 1:100, Hybridoma Bank) washed and incubated with secondary antibodies coupled to Alexa dyes and rhodamine-phalloidin or phalloidin-Alexa-660 (all from Molecular Probes). After washing, samples were mounted in Vectashield containing DAPI. Images were acquired with a Zeiss LSM 780 confocal microscope and processed with Fiji (Schindelin et al., 2012) and Photoshop (Adobe).

### Live imaging

SPIM imaging of embryos: De-chorionated embryos of the appropriate age were embedded in 1 % low melting point agarose and mounted into a glass capillary. Fluorescent microspheres (FY050 Estapor microspheres, Merck Millipore; 1:4000) were included in the embedding medium for multi-view registration. The embryos were imaged using the Zeiss Lightsheet Z.1 with a Zeiss 20x/1.0 water-immersion Plan Apochromat objective lens with 0.8x zoom at 25 °C using 488 nm laser set at 4 mW. Five views were imaged using dual-sided illumination with Zeiss 10x/0.2 illumination lenses. A mean fusion was applied to fuse both illumination sides after acquisition using the ZEN software (Zeiss). The views were acquired at 72° angles with a stack size of 130 μm and a step size of 1.5 μm. Exposure time were 30 ms per slice. Each slice consists of 1920 × 1200 pixels with a pixel size of 0.29 μm and a bit depth of 16 bits. The light sheet thickness was 4 μm at the center of the field of view. The embryos were imaged from onset of expression of the fosmids transgenes (determined empirically) until late embryogenesis with a time resolution of 15 min. Multi-view processing of the dataset was carried out using the Fiji plugin for multiview reconstruction (Preibisch et al., 2009; Schmied et al., 2014), which was executed on a high performance computing cluster (Schmied et al., 2015). The multi-view reconstruction was followed by multi-view deconvolution (Preibisch et al., 2014), for which the images were down sampled by a factor of two. Movies were extracted via the Fiji plugin BigDataViewer (Pietzsch et al., 2015).

The Gsb-GFP fTRG line was crossed with the H2Av-mRFPruby line (Fischer et al., 2004; Preibisch et al., 2014), the embryos of this cross were imaged using a 40x/1.0 water immersion Plan Apochromat lens from Zeiss with 1x zoom at 25 °C at 17.5 mW of the 488 nm laser and 4 mW of the 561 nm laser. A single angle with dual sided illumination was imaged. The stack size was 82.15 μm with a step size of 0.53 μm. Exposure time was 30 ms per slice. Each slice consisted of 1920 × 1920 pixels with a pixel size of 120 nm and a bit depth of 16-bit. The light sheet thickness was 3.21 μm at the center of the field of view. The embryos were imaged from early blastoderm onwards until late embryogenesis focusing on the head with a time resolution of 7 min.

Imaging of pupae: Staging and live imaging of the pupae were performed at 27 °C. Live imaging of pupae at the appropriate stage was done as described previously (Weitkunat and Schnorrer, 2014). Briefly, the staged pupa was cleaned with a brush and a small observation window was cut into the pupal case with sharp forceps. The pupa was mounted on a custom-made slide and the opening was covered with a small drop of 50 % glycerol and a cover slip. Z-stacks of either single time points or longterm time lapse movies were acquired using either a spinning disc confocal microscope (Zeiss, Visitron) or a two-photon microscope (LaVision), both equipped with heated stages.

### Proteomics

Per sample about hundred pupae or adult flies were snap-frozen in liquid nitrogen and ground to a powder. The powder was re-suspended and further processed as described in the quantitative BAC-GFP interactomics protocol (Hubner et al., 2010). In brief, 800 μl of lysate per sample were cleared by centrifugation. The cleared lysate was mixed with magnetic beads pre-coupled to anti-GFP antibodies and run over magnetic micro-columns (both Miltenyi Biotec). Columns were washed, and samples subjected to in-column tryptic digestion for 30 min. Eluates were collected and digestion continued overnight, followed by desalting and storage on StageTips. Eluted peptides were analysed with an Orbitrap mass spectrometer (Thermo Fisher). Raw data were analysed in MaxQuant version 1.4.3.22 (Cox and Mann, 2008) using the MaxLFQ algorithm for label-free quantification (Cox et al., 2014). Interacting proteins were identified by the similarity of their intensity profiles to the respective baits (Keilhauer et al., 2014). Heat maps were plotted in the Perseus module of the MaxQuant software suite.

## Author contributions

D.S., S.H., E.V. and M.S performed the liquid culture recombineering experiments. M.S., S.J., P.K. and S.S. analysed the NGS data and contributed to the TransgeneOme database.

B. S., N.P., K.F. and F.S performed the transgenesis of about half the fTRG lines. V.K.J.V., R.T.K., K.A., M.R. and K.V performed the transgenesis of the other half of the fTRG lines.

C. B. performed most of the genetic rescue experiments.

H.J. performed and analysed all experiments in ovaries.

C.S. collected and analysed SPIM in toto images of embryos.

C.B. and F.S. performed and analysed all imaging data in pupae and adults.

M.Y.H and M.M. performed the proteomics analysis and analysed the data.

V.H. contributed analysis of Gsb-GFP cell tracking.

R.K.E. constructed the tagging cassettes.

I.R.S.F. performed initial proof of principle tagging experiments.

P.T., F.S., M.S. and E.K. initiated the collaborative project and obtained dedicated funding.

P.T. and F.S. conceived and coordinated the study and co-wrote the manuscript with input from the other authors.

## Acknowledgements

Stocks obtained from the Bloomington *Drosophila* Stock Center (NIH P40OD018537) were used in this study. We thank Anne Ephrussi for sharing of fly lines. We thank Franziska Friedrich for drawing the ovariole scheme used for Figure 4A. We thank Andreas Dahl from the Deep Sequencing Group at CRTD/BIOTECH, Dresden for the NGS library preparation and sequencing. We also thank light microscopy facility and the computer department for assistance with imaging and data processing. We are grateful to Sandra Lemke, Aynur Kaya-Copur and Xu Zhang for help with the genetic rescue experiments and to Caroline Sonsteby for assistance during some of the pupal imaging experiments. We thank the entire Schnorrer lab for helpful comments on this manuscript. We are particularly grateful to Reinhard Fässler and Herbert Jäckle for continuous support of this work. This work was funded by the EMBO Young Investigator Program (F. S.) the European Research Council under the European Union’s Seventh Framework Programme (FP/2007-2013)/ERC Grant 310939 (F. S.), the European Research Council under the European Union’s Seventh Framework Programme (FP/2007-2013)/ERC Grant 260746 (P.T.), European Commission GENCODYS (P.T. and H.J.), Human Frontier Science Program (HFSP) RGY0093/2012 (P.T. and C. S.) and the Max Planck Society (P.T., F.S., M.S., M.M. and E.K.).

---

**Figure 2 - Supplement 1:**
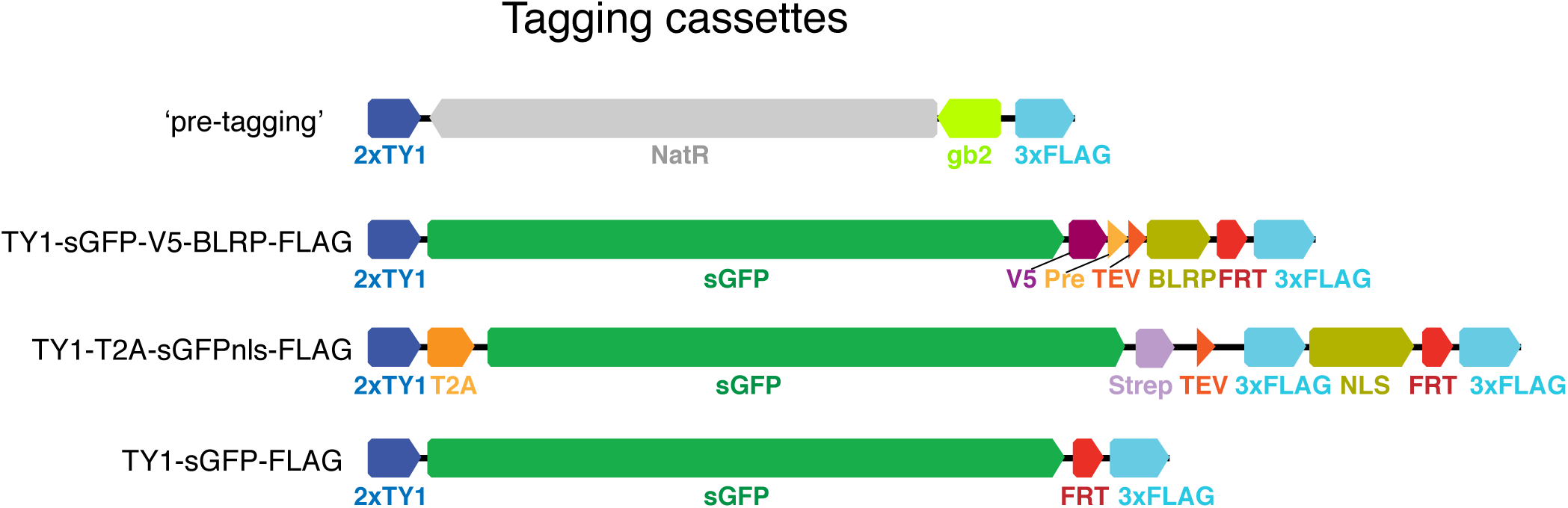
Tagging cassettes. Tags tested and used in this study. Shown is the form of the tagging cassette after insertion into target site and flip-out of the counter selection sequences (rpsl-neo) leaving behind the FRT sequence colored red.

**Figure 4 - Supplement 1:**
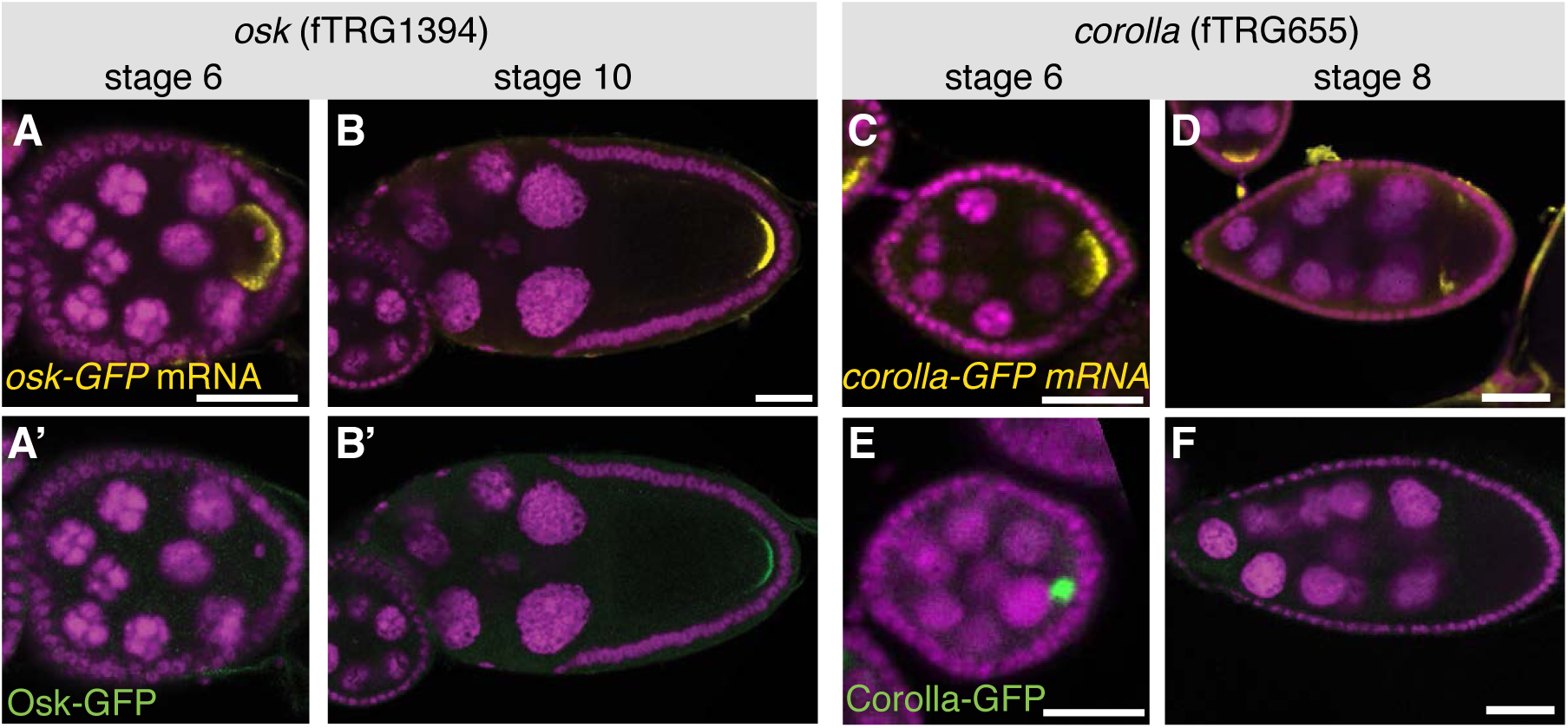
Posttranscriptional regulation of protein expression during oogenesis. (**A**, **B**) *osk-GFP* mRNA visualised by an anti-GFP labelled RNA probe (yellow, DAPI in magenta) at stage 6 and stage 10 of oogenesis. (**A**’, **B**’) Osk-GFP protein visualised by anti-GFP antibody (green, DAPI in magenta) at stage 6 and stage 10. Note that Osk-GFP protein is not detectable at stage 6. (**C, D**) *corolla-GFP* mRNA (yellow, DAPI in magenta) at stage 6 and stage 8. (**E, F**) Corolla-GFP protein (green, DAPI in magenta) at stage 6 and stage 8. Note that Corolla-GFP protein in only detectable at stage 6 but not stage 8. Scale bars indicate 30 μm.

**Figure 5 - Supplement 1:**
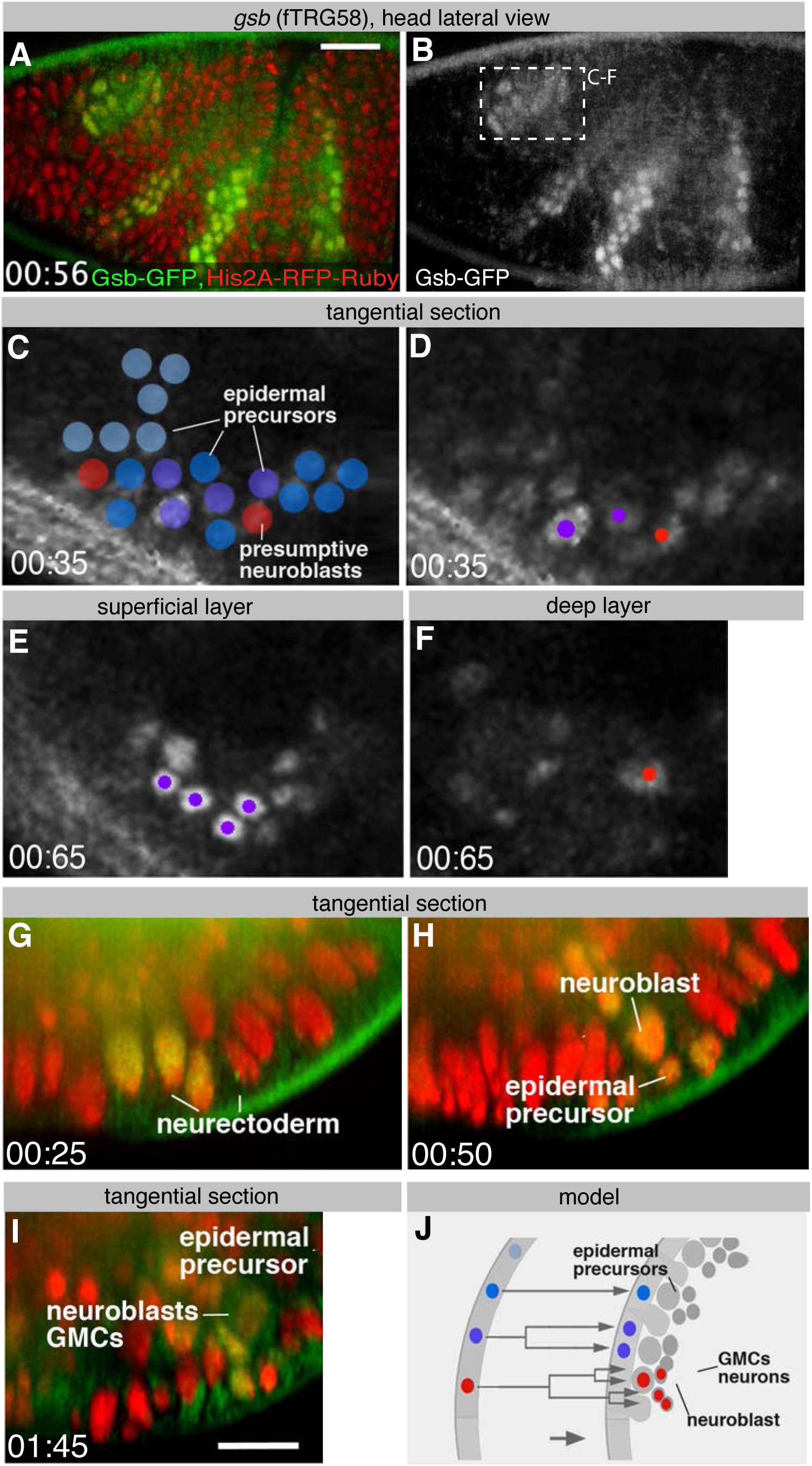
Live Gsb-GFP imaging during embryogenesis with SPIM. (**A**, **B**) Lateral view of a live stage 10 embryo expressing Gsb-GFP (green in A) and Histone2A-mRFPruby (red in A); anterior is to the left (see **Supplementary Movie 2**). (**C** - **F**) Reconstruction of *gsb*-positive deuterocerebral proneural domain (from a similar movie and position as boxed in B). Fate of cells is symbolized by different colours (blue: epidermal precursor undergoing no further mitosis; purple: epidermal precursor undergoing one mitosis; red: neuroblasts). (C) and (D) show optical section through neurectoderm of stage 10 embryo prior to neuroblast delamination; (E) and (F) 30 minutes later, after neuroblasts have delaminated (stage 11), with a superficial optical section of surface ectoderm (E), and deep section of neuroblast layer (F). (**G** - **I**) Optical cross sections of similar live embryo as in (A) expressing GFP-tagged Gsb (green) and Histone-2A-mRFPruby (red) at stage 10 (G), early 11 (H) and late 11 (I) showing neuroblasts delaminating from Gsb-GFP domain. **J**: Schematic cross section of stage 10 (left) and stage 11 (right) ectoderm illustrating fate of cells forming part of Gsb-positive pro-neural domain. Scale bars indicate 25 μm (A, B) and 10 μm (G - I).

**Figure 5 - Supplement 2:**
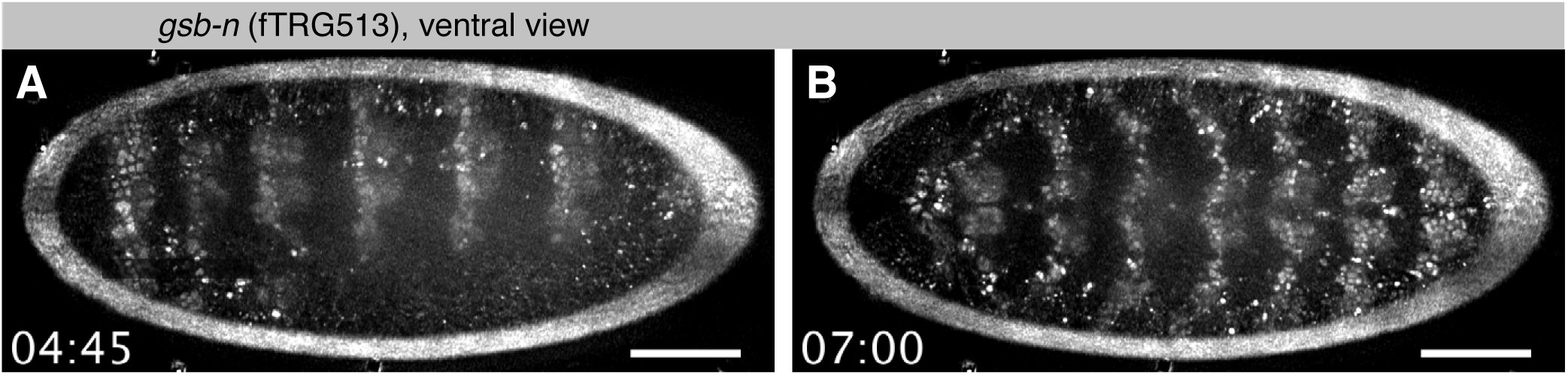
Live Gsb-n-GFP imaging during embryogenesis with SPIM. (**A, B**). Ventral view of Gsb-n-GFP expression of a stage 12 embryo during germband retraction (A) and stage 14 during head involution (B). Note that Gsb-n-GFP remains expressed in neuronal precursors during stage 14 (**Supplementary Movie 3**). Scale bars indicate 50 μm.

**Figure 6 - Supplement 1:**
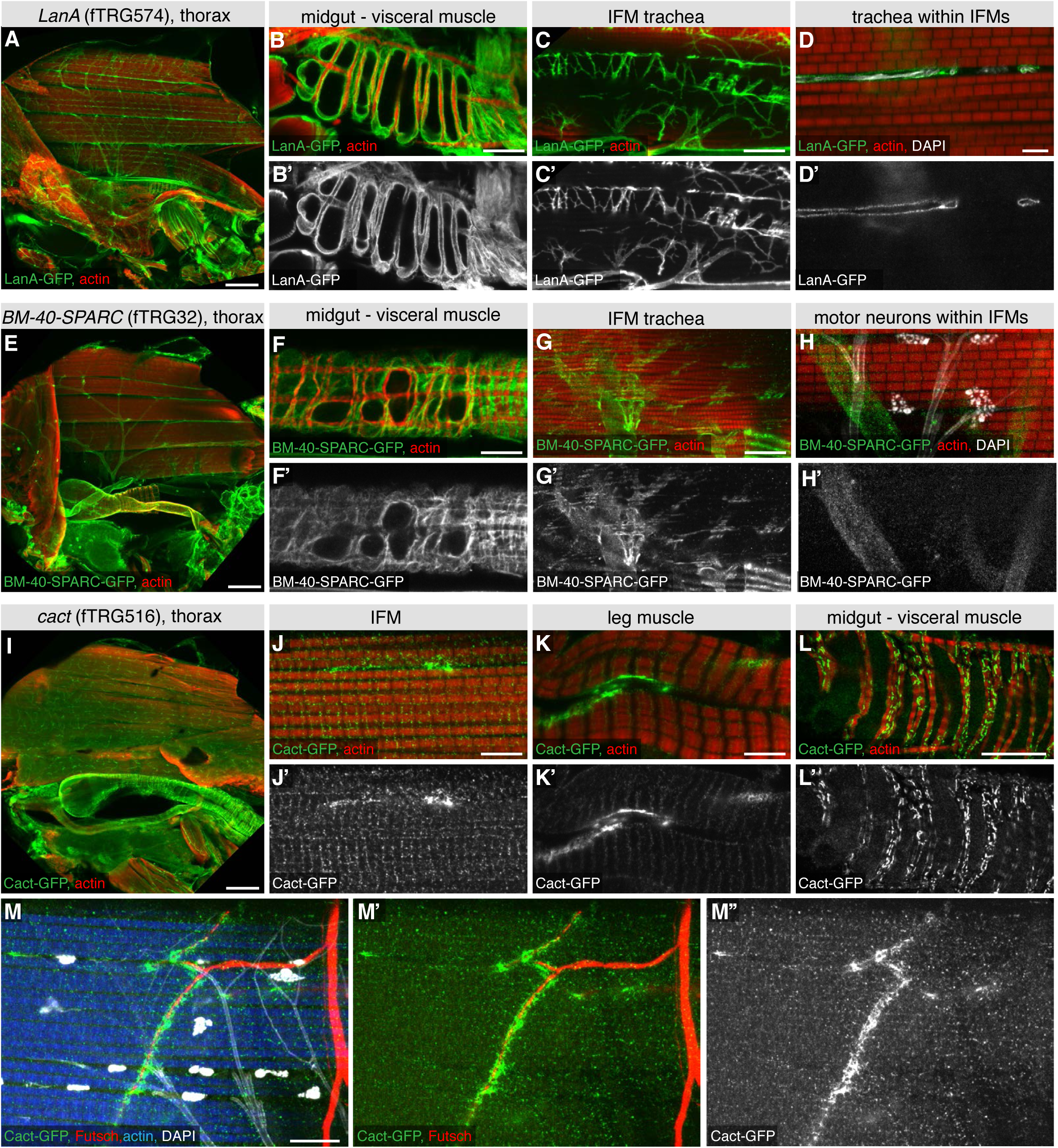
ECM and synaptic markers of the adult thorax. (**A** - **H**) Antibody stainings of the adult thorax with anti-GFP antibody (green or white in the single colour images) and phalloidin (red). LanA-GFP and BM-40-SPARC-GFP labels the ECM around motor neurons, visceral muscle and trachea. Note that thin trachea within the IFMs (marked by UV auto-fluorescence in white) are surrounded by LanA-GFP but not BM-40-SPARC-GFP (**D**, **H**). (**I** - **M**) Cact-GFP (green) shows a distinct localisation in IFMs, leg and visceral muscle reminiscent of a neuromuscular junction pattern. Note the partial co-localisation with the motor neuron marker Futsch (in red, **M, M’**), whereas no co-localisation with trachea in IFMs (in white, **M**). Scale bars indicate 100 μm (A, E, I), 20 μm (B, C, F, G, L), 10 μm (M) and 5 μm (D, H, J, K).

**Figure 7 - Supplement 1:**
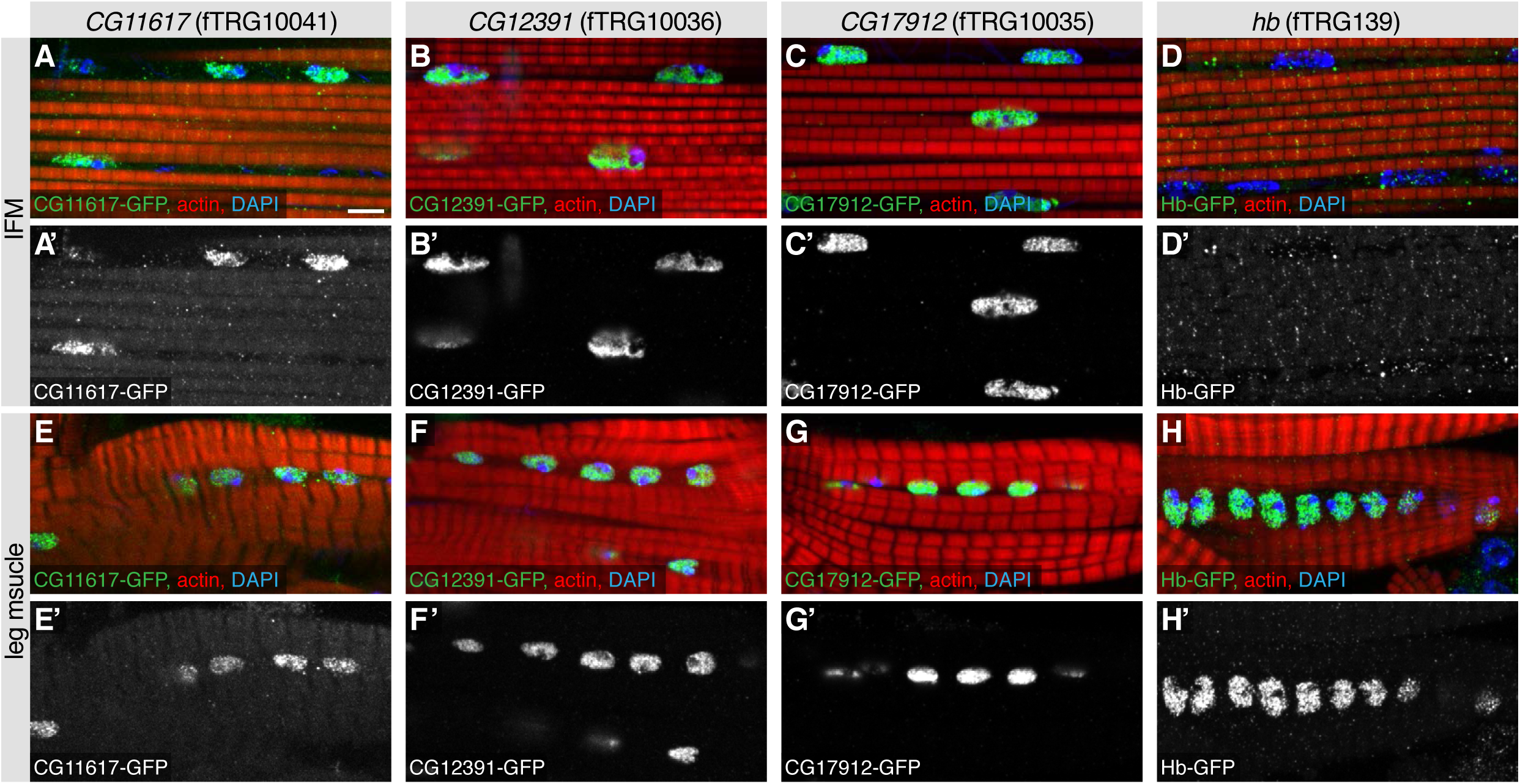
Nuclear localisations in adult flight muscles. Antibody stainings of the adult thorax with anti-GFP antibody (green or white in the single colour images), phalloidin (red) and DAPI (blue). (**A** - **H**) CG11617-GFP (**A**, **E**), CG12391-GFP (**B**, **F**) and CG17912 (**C**, **G**) are localised to the nuclei of IFMs and leg muscles, whereas Hb-GFP is only found in leg muscle nuclei (**H**) and not detectable in IFM nuclei (**D**). Scale bars indicate 5 μm.

**Figure 7 - Supplement 2:**
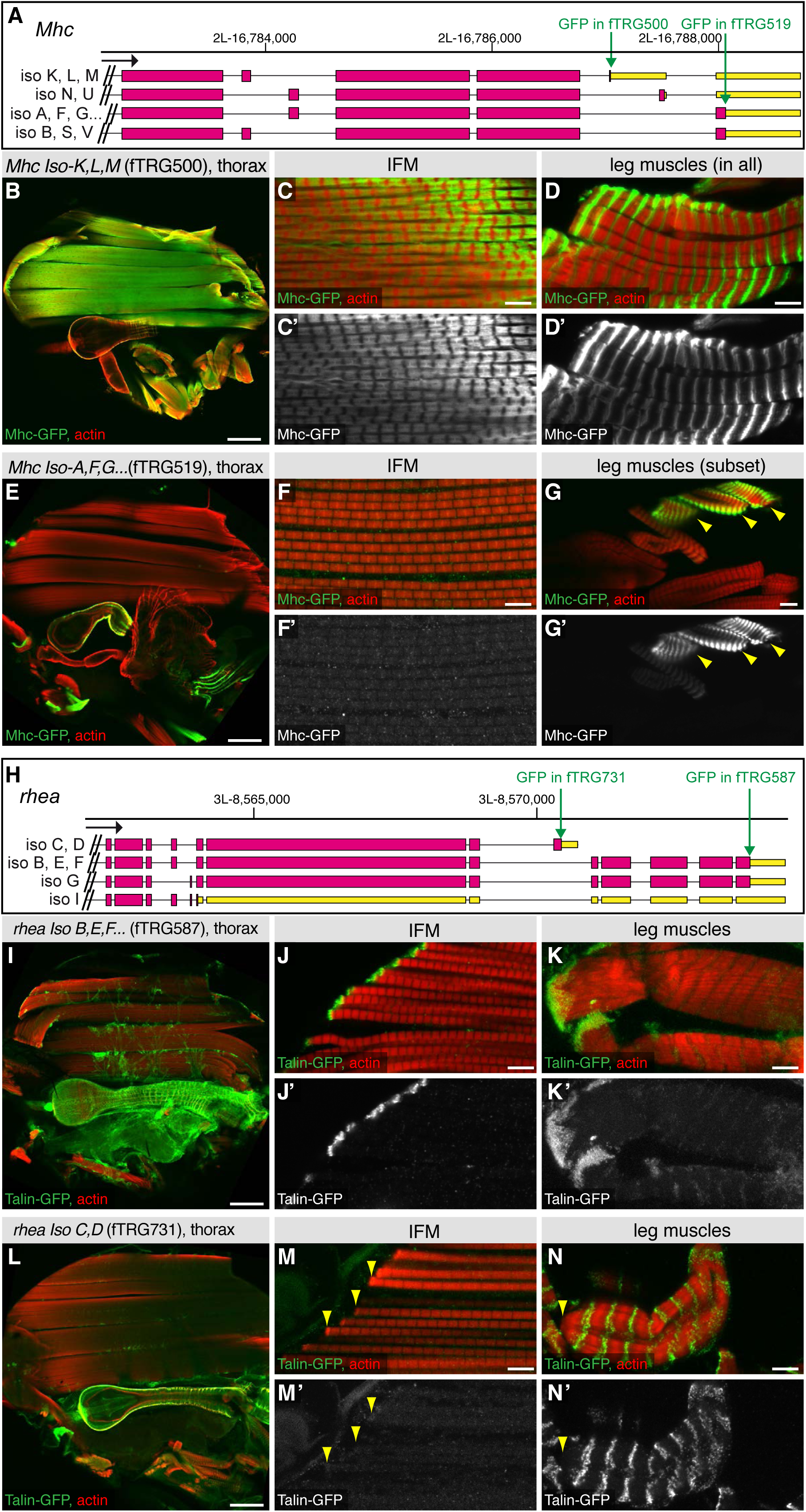
Alternative splicing into alternative C-termini. Antibody stainings of the adult thorax with anti-GFP antibody (green or white in the single colour images) and phalloidin (red). (**A, H**) Gene models of the 3’ end of *Mhc* (**A**) and *rhea* (**H**) listing the predicted isoforms; coding exons are shown in pink, 3’UTRs in yellow boxes. The positions of the GFP tag insertions are marked by green arrows. (**B** - **G**) The shorter Mhc-GFP isoforms (Iso K, L, M) are expressed in IFMs and all leg muscles (**B** - **D**), whereas the slightly longer Mhc-GFP isoforms (Iso A, F, G etc.) are not detectable in IFMs but present in visceral muscles and a subset of leg muscles (**E** - **G**). (**I** - **N**) The long Talin-GFP isoforms (*rhea* Iso B, E, F, G) localise to muscle-tendon attachment sites in IFMs (**J**) and leg muscles (**K**), whereas the shorter Talin-GFP isoforms (*rhea* Iso C, D) are not detectable at muscle-tendon attachment sites in IFMs (**M**, arrowheads) and leg muscles (**N**, arrowhead), however do localise to costamers of leg muscles (**N**). Scale bars indicate 100 μm (B, E, I, L), 10μm (G) and 5 μm (C, D, F, J, K, M, N).

**Figure 8 - Supplement 1:**
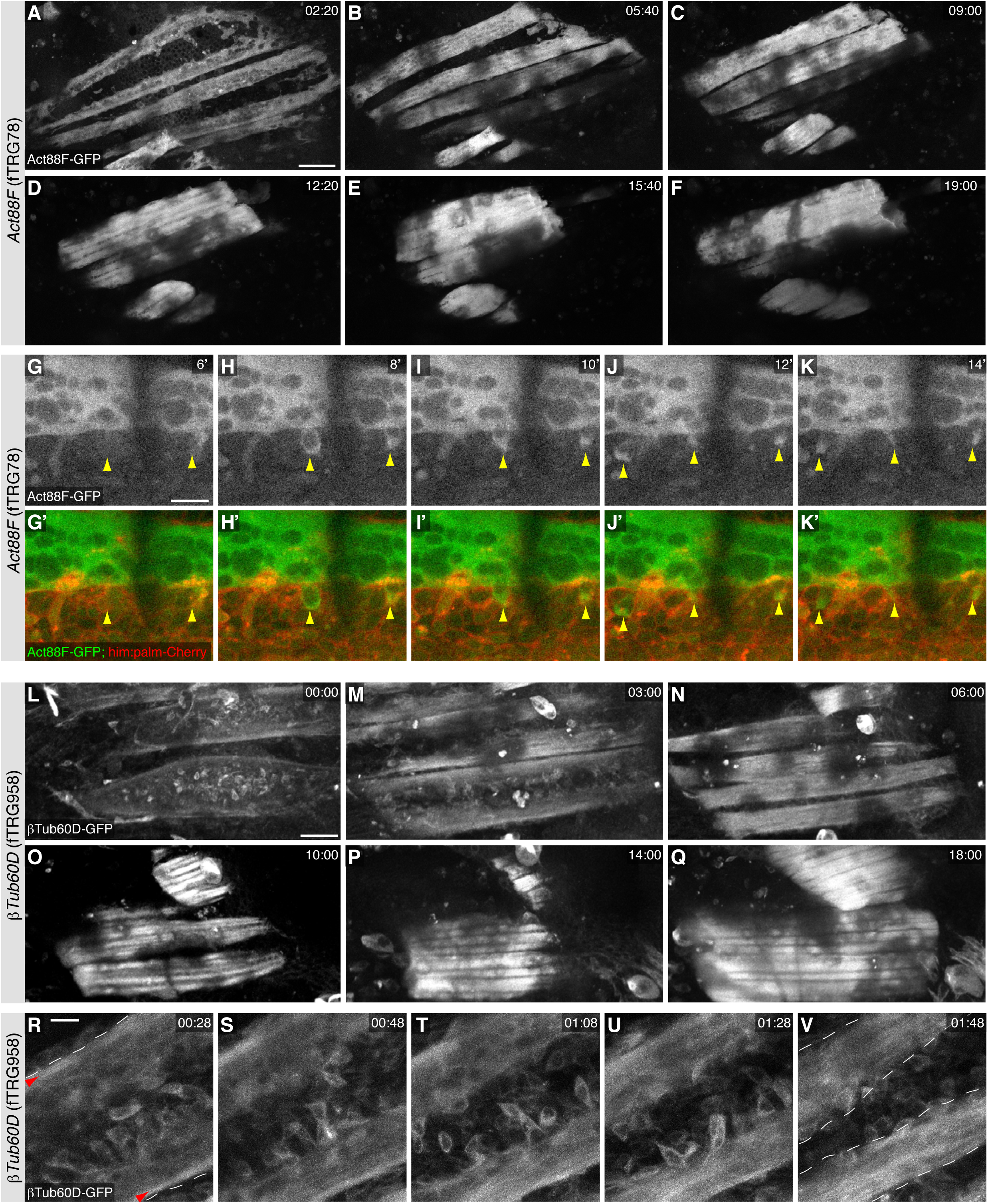
Live imaging during pupal development. (**A** - **F**) Stills from live two-photon imaging of an intact 14h APF pupa expressing Act88F-GFP strongly labelling the developing IFMs (see **Supplementary Movie 4**). (**G** - **K**) Stills from a two-colour spinning disc movie expressing Act88F-GFP (green) and *him-*GAL4, UAS-palm-Cherry (red) labelling the myoblasts (see **Supplementary Movie 5**). Note the sudden green label of single myoblasts after fusion had occurred (yellow arrowheads, see **Supplementary Movie 5**). (**L** - **Q**) Stills from two-photon movie of an intact 14h APF pupa expressing βTub-60D-GFP in fusing myoblasts and developing myofibers (See **Supplementary Movie 6**). (**R - V**) Still from a high resolution two-photon movie of an intact 16h APF pupa expressing βTub-60D-GFP. Single myoblast during fusion can be resolved (See **Supplementary Movie 7**). Strong microtubule bundles (red arrow heads) are visible close to the edges of the splitting myotube (white dashed lines, **R**); splitting is complete in (**V**). Scale bars indicate 50 μm (A - F, L - Q) and 10 μm (G - K) and (R - V).

**Supplementary Table 1:** TransgeneOme constructs. Construct and clone names, tag locations, as well the sequencing validation data are listed for all TransgeneOme constructs generated. Sheet 1 lists the sGFP TransgeneOme (TY1-sGFP-V5-BLRP-FLAG tag, NGS sequenced), sheet 2 the TY1-sGFP-FLAG pilot set clones (junctions Sanger sequenced, only exact matches are counted as verified), sheet 3 the TY1-T2A-sGFPnls-FLAG pilot set clones (entire tag Sanger sequenced) and sheet 4 the TY1-sGFP-V5-BLRP-FLAG pilot set clones (entire tag Sanger sequenced). Sheet 5 summarises all verified tagged genes in these sets.

**Supplementary Table 2:** Transgenic FlyFos (fTRG) lines. Table listing all 880 transgenic FlyFos (fTRG) lines, with fTRG numbers, construct and clones names, as well as nature of the tag and the used landing site. The second sheet compares the genes tagged by the fTRG lines to the available GFP gene trap lines. 765 genes are only found in the TransgeneOme resource.

**Supplementary Table 3:**
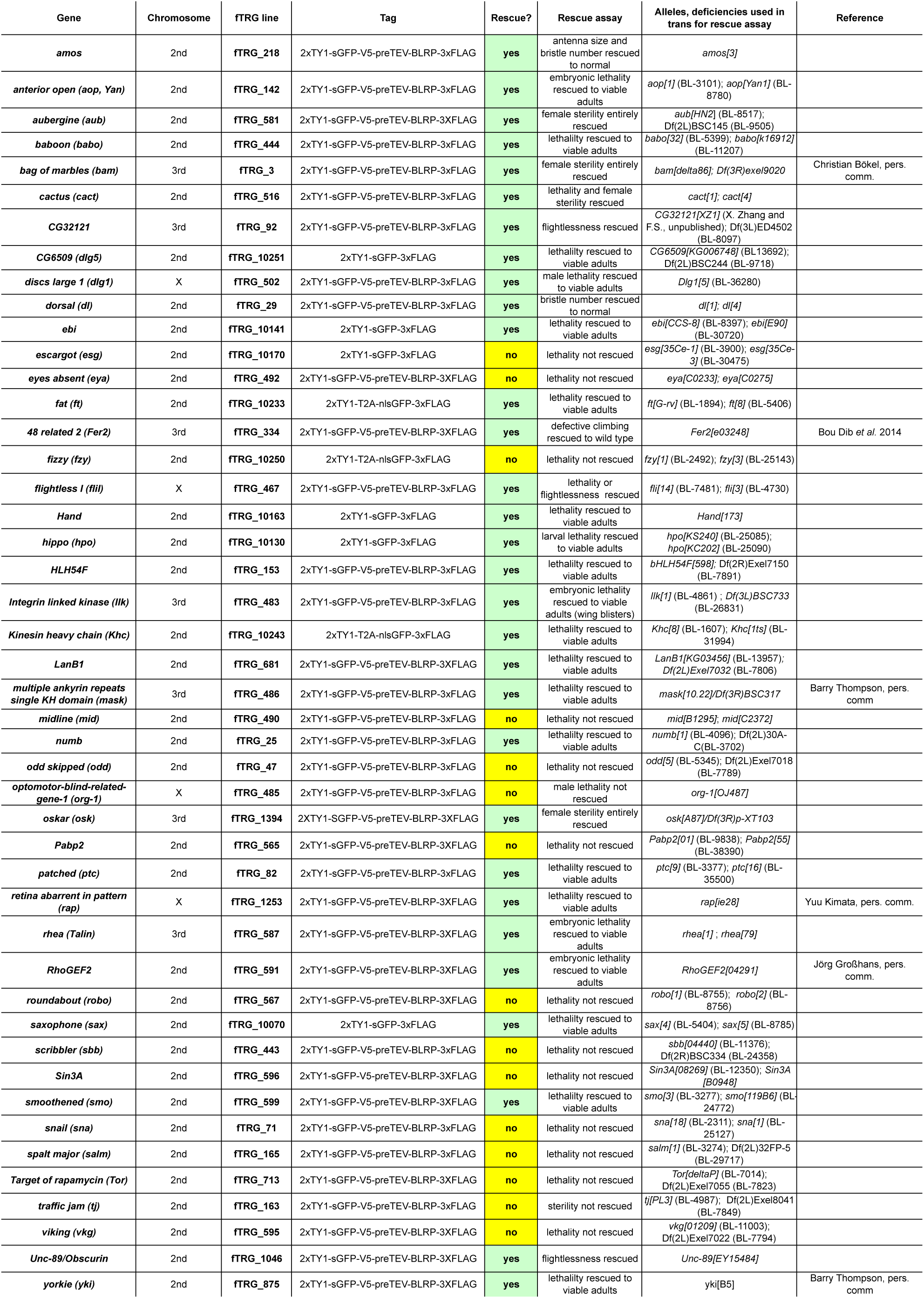
Genetic rescue of the fTRG lines. Table listing fTRG lines and respective genetic alleles as well as rescue assays that were used to assess the functionality of the fTRG lines. Note that about two-thirds of the lines fully rescue the mutant phenotypes (marked green).

**Supplementary Table 4:**
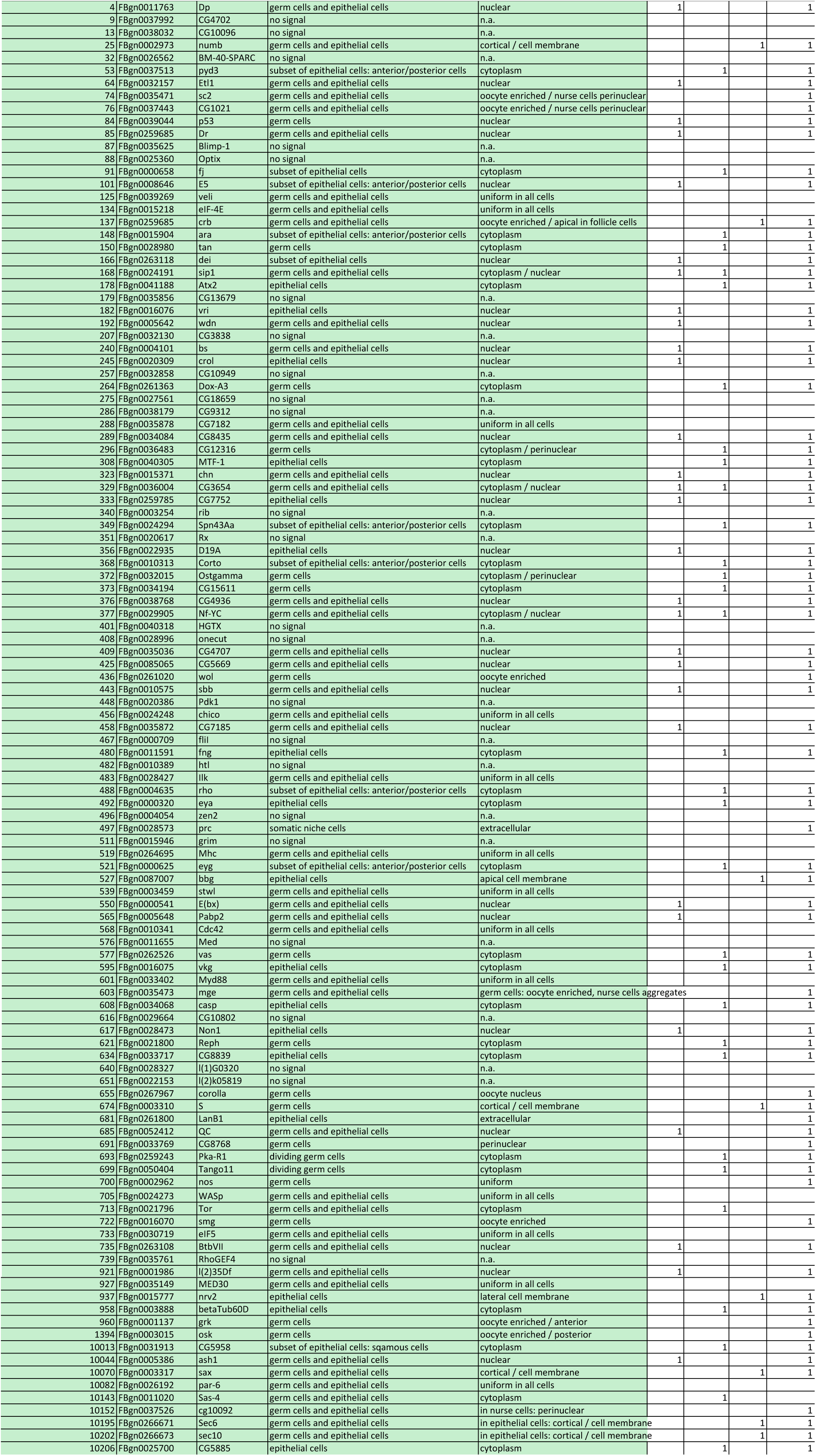
FlyFos (fTRG) expression in ovaries. Table listing the expression patterns for 115 fTRG lines in ovaries. Expression was detected in 94 lines by anti-GFP antibody stainings. Cell type specific expression and subcellular localisations were monitored for these lines.

**Supplementary Table 5:**
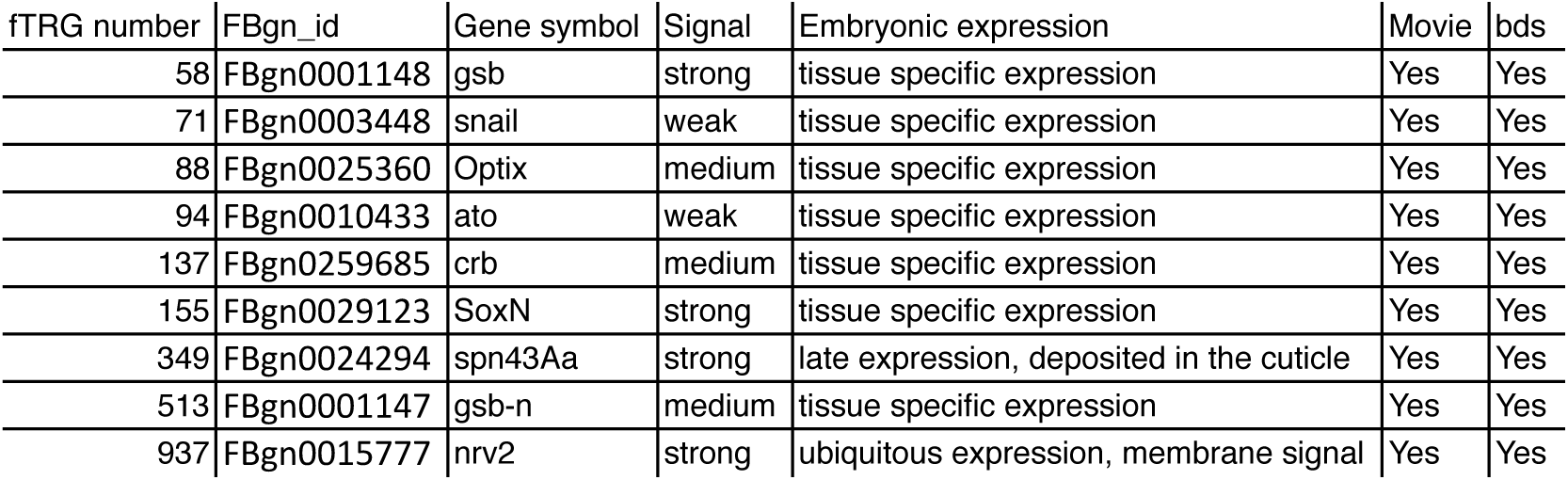
*in toto* SPIM imaging of FlyFos (fTRG) lines in the embryo. Table listing the fTRG lines that were imaged in the embryo using Zeiss Lightsheet Z.1 from multiple angles over time. *nrv2, gsb* and *gsb-n* are discussed in the text. For the remaining lines we list broad categorisation of the expression detected by SPIM imaging.

| fTRG number | FBgn_id | Gene symbol | Signal | Embryonic expression | Movie | bds |
| --- | --- | --- | --- | --- | --- | --- |
| 58 | FBgn0001148 | gsb | strong | tissue specific expression | Yes | Yes |
| 71 | FBgn0003448 | snail | weak | tissue specific expression | Yes | Yes |
| 88 | FBgn0025360 | Optix | medium | tissue specific expression | Yes | Yes |
| 94 | FBgn0010433 | ato | weak | tissue specific expression | Yes | Yes |
| 137 | FBgn0259685 | crb | medium | tissue specific expression | Yes | Yes |
| 155 | FBgn0029123 | SoxN | strong | tissue specific expression | Yes | Yes |
| 349 | FBgn0024294 | spn43Aa | strong | late expression, deposited in the cuticle | Yes | Yes |
| 513 | FBgn0001147 | gsb-n | medium | tissue specific expression | Yes | Yes |
| 937 | FBgn0015777 | nrv2 | strong | ubiquitous expression, membrane signal | Yes | Yes |

**Supplementary Table 6:**
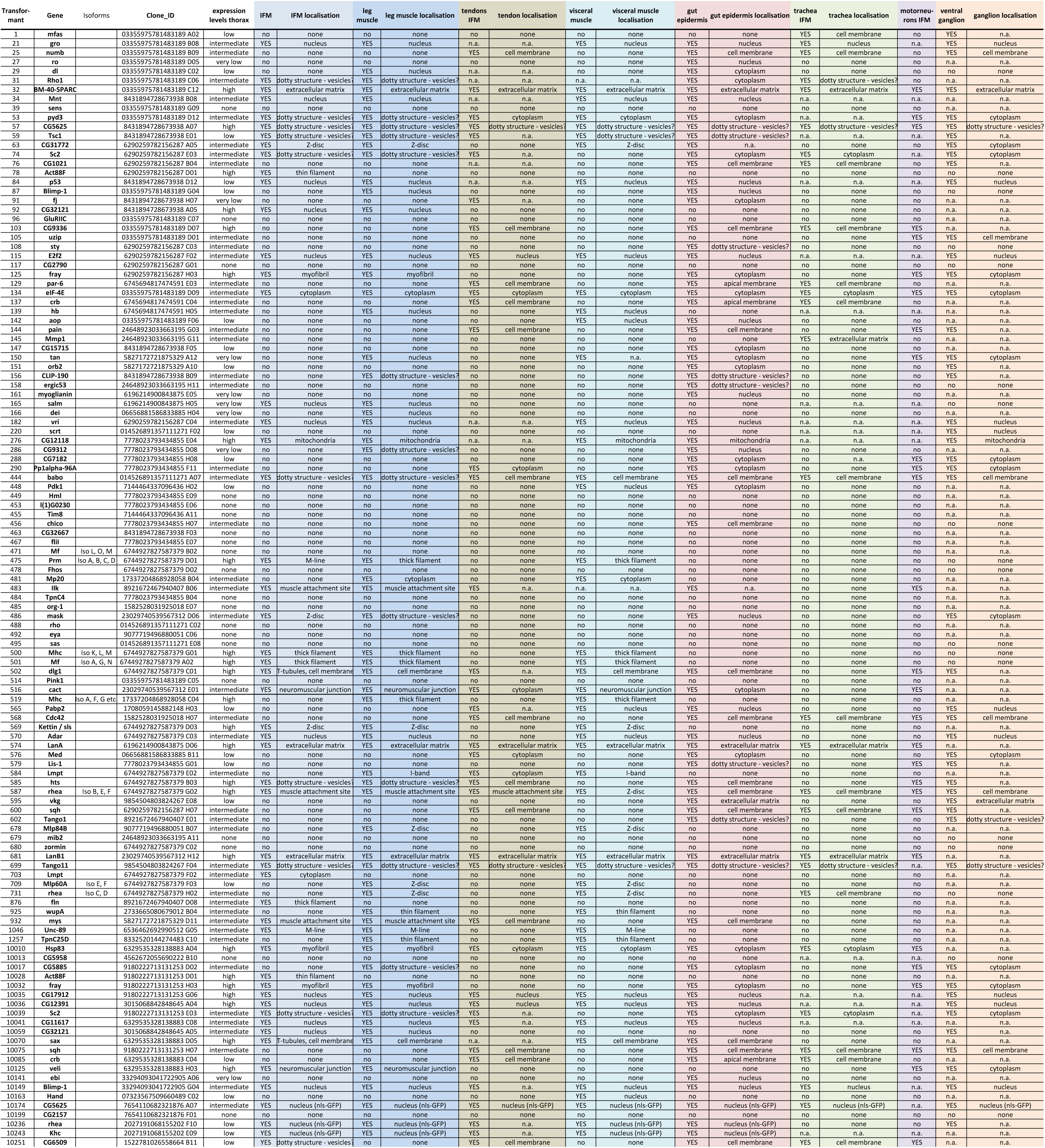
FlyFos (fTRG) expression in the adult thorax. Table listing the expression pattern for 121 fTRG lines in adult thoraces. Expression was detected in101 lines by anti-GFP antibody stainings. Cell type specific expression and subcellular localisations were monitored for these lines.

**Supplementary Table 7:**
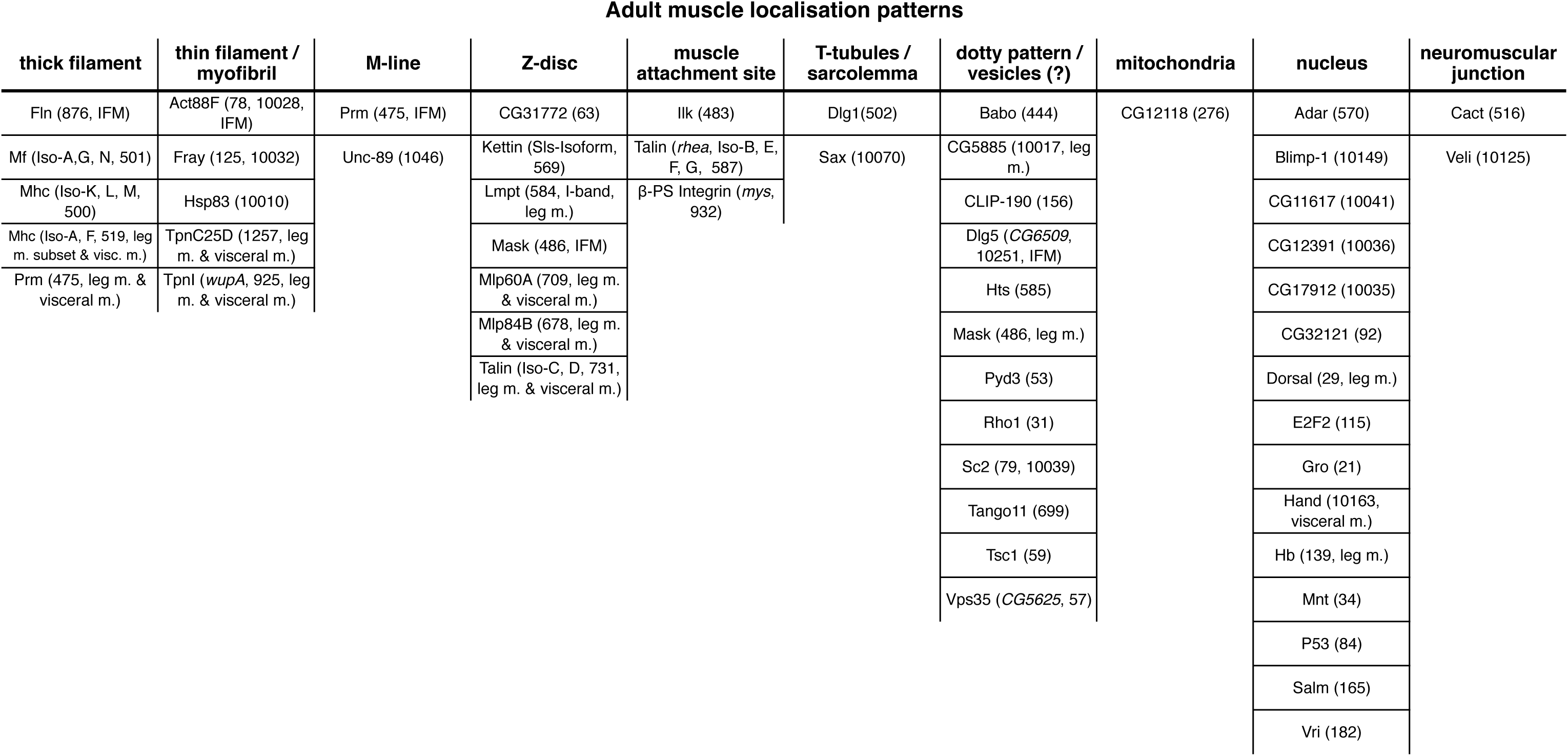
Summary of adult muscle patterns of FlyFos (fTRG) lines. 54 detected adult muscle localisation patterns (flight muscle, leg muscle and visceral muscle) from Supplementary Table 6 are summarised.

**Supplementary Table 8:** Proteomics quantification. Quantitative mass spectrometry values of all detected protein obtained with the MaxQuant software suite for all the GFP-enrichment experiments are listed.

**Supplementary Movie 1**

Multi-view SPIM movie of a stage 12 Nrv2-GFP expressing embryo. A stack was acquired every 15 minutes, lateral, dorsal, ventral and transverse views of the same time points are displayed. From stage 11 onwards Nrv2-GFP is present ubiquitously in the plasma membrane. Later its expression increases in the CNS, particularly in the neuropil and the motor neurons. Movie plays with 7 frames per second. Time is given in hh:mm. Scale bar indicates 50 μm.

**Supplementary Movie 2**

Lateral head section from a SPIM movie of a stage 10 Gsb-GFP (green, white in the top movie), Histone-2A-mRFPruby (red) embryo. A stack was acquired every 7 minutes. The segmentally re-iterated stripe-like *gsb* expression domain in the head neuroectoderm is visible. Later, *gsb* is expressed in ganglion mother cells and nerve cells that are the progeny of *gsb* expressing neuroblasts. Movie plays with 7 frames per second. Time is given in hh:mm. Scale bar indicates 50 μm.

**Supplementary Movie 3**

Ventral view of a SPIM movie of a stage 6 Gsb-n-GFP embryo. A stack was acquired every 15 minutes. Gsb-n-GFP is only detectable at the end of germ-band extension. During germ-band retraction it is expressed in characteristic L-shaped expression domains in the hemi-segments of the trunk. In the late stage embryo Gsb-n-GFP is present in the neurons of the shortening ventral nerve cord. Movie plays with 7 frames per second. Time is given in hh:mm. Scale bar indicates 50 μm.

**Supplementary Movie 4**

Z-projection of a two-photon movie of an about 14h APF pupa expressing Act88F-GFP. A stack was acquired every 20 min for 19 h. Expression of Act88F-GFP increases in the indirect flight muscles dramatically, thus contrast was reduced several times in course of the movie to avoid over-exposure. Movie plays with 5 frames per second. Time is given in hh:mm.

**Supplementary Movie 5**

Single plane of a spinning disc confocal movie of an about 14 h APF old pupa expressing Act88F-GFP (green) in the flight muscle myotubes and *him*-GAL4; UAS-palm-Cherry in the myoblasts. An image stack was acquired every two minutes. Note the newly fused myoblasts acquired the GFP label within a single time interval (highlighted by green arrows). Movie plays with 5 frames per second. Time is given in minutes.

**Supplementary Movie 6**

Z-projection of a two-photon movie of an about 14h APF pupa expressing βTub60D-GFP. A stack was acquired every 20 min for 25 h. Note the high expression of βTub60D-GFP in fusing myoblasts and the thick microtubules bundles in the developing flight muscles. Hair cells of the developing sensory organs also show strong expression, however move out of the Z-stack over time. Movie plays with 5 frames per second. Time is given in hh:mm.

**Supplementary Movie 7**

Single plane of a two-photon movie of an about 16 h APF old pupa expressing βTub60D-GFP in myoblasts and the forming flight muscle myotubes. An image stack was acquired every two minutes for more than 3 h. Note that single myoblasts can be followed during fusion. Most myoblasts fuse in the center of the myotube, which gradually splits into two myotubes. Movie plays with 5 frames per second. Time is given in hh:mm.

## Footnotes

1 Examine raw data in BigDataViewer (Fiji -> Plugins -> BigDataViewer -> Browse BigDataServer and http://bds.mpi-cbg.de:8087

## References

Adams, M.D., Celniker, S.E., Holt, R.A., Evans, C.A., Gocayne, J.D., Amanatides, P.G., Scherer, S.E., Li, P.W., Hoskins, R.A., Galle, R.F., et al. (2000). The genome sequence of Drosophila melanogaster. Science 287, 2185–2195.

Baena-Lopez, L.A., Alexandre, C., Mitchell, A., Pasakarnis, L., and Vincent, J.P. (2013). Accelerated homologous recombination and subsequent genome modification in Drosophila. Development 140, 4818–4825.

Bard, F., Casano, L., Mallabiabarrena, A., Wallace, E., Saito, K., Kitayama, H., Guizzunti, G., Hu, Y., Wendler, F., DasGupta, R., et al. (2006). Functional genomics reveals genes involved in protein secretion and Golgi organization. Nature 439, 604–607.

Bischof, J., Bjorklund, M., Furger, E., Schertel, C., Taipale, J., and Basler, K. (2013). A versatile platform for creating a comprehensive UAS-ORFeome library in Drosophila. Development 140, 2434–2442.

Bolatto, C., Chifflet, S., Megighian, A., and Cantera, R. (2003). Synaptic activity modifies the levels of Dorsal and Cactus at the neuromuscular junction of Drosophila. J. Neurobiol. 54, 525–536.

Bonnay, F., and Cohen-Berros, E. (2013). big bang gene modulates gut immune tolerance in Drosophila.

Bulgakova, N.A., and Knust, E. (2009). The Crumbs complex: from epithelial-cell polarity to retinal degeneration. Journal of Cell Science 122, 2587–2596.

Buszczak, M., Paterno, S., Lighthouse, D., Bachman, J., Planck, J., Owen, S., Skora, A.D., Nystul, T.G., Ohlstein, B., Allen, A., et al. (2007). The carnegie protein trap library: a versatile tool for Drosophila developmental studies. Genetics 175, 1505–1531.

Clark, K.A., Bland, J.M., and Beckerle, M.C. (2007). The Drosophila muscle LIM protein, Mlp84B, cooperates with D-titin to maintain muscle structural integrity. Journal of Cell Science 120, 2066–2077.

Collins, K.A., Unruh, J.R., Slaughter, B.D., and Yu, Z. (2014). Corolla is a novel protein that contributes to the architecture of the synaptonemal complex of Drosophila. ….

Cox, J., and Mann, M. (2008). MaxQuant enables high peptide identification rates, individualized p.p.b.-range mass accuracies and proteome-wide protein quantification. Nature Biotechnology 26, 1367–1372.

Cox, J., Hein, M.Y., Luber, C.A., Paron, I., Nagaraj, N., and Mann, M. (2014). Accurate proteome-wide label-free quantification by delayed normalization and maximal peptide ratio extraction, termed MaxLFQ. Mol. Cell Proteomics 13, 2513–2526.

de Celis, J. (1999). Regulation of the spalt/spalt-related gene complex and its function during sensory organ development in the Drosophila thorax. Development 126, 2653.

Deal, R.B., and Henikoff, S. (2010). A simple method for gene expression and chromatin profiling of individual cell types within a tissue. Developmental Cell 18, 1030–1040.

Dietzl, G., Chen, D., Schnorrer, F., Su, K.-C., Barinova, Y., Fellner, M., Gasser, B., Kinsey, K., Oppel, S., Scheiblauer, S., et al. (2007). A genome-wide transgenic RNAi library for conditional gene inactivation in Drosophila. Nature 448, 151–156.

Dougherty, G.W., Chopp, T., Qi, S.-M., and Cutler, M.L. (2005). The Ras suppressor Rsu-1 binds to the LIM 5 domain of the adaptor protein PINCH1 and participates in adhesion-related functions. Experimental Cell Research 306, 168–179.

Dunst, S., Kazimiers, T., Zadow, von, F., Jambor, H., Sagner, A., Brankatschk, B., Mahmoud, A., Spannl, S., Tomancak, P., Eaton, S., et al. (2015). Endogenously tagged rab proteins: a resource to study membrane trafficking in Drosophila. Developmental Cell 33, 351–365.

Ejsmont, R.K., Sarov, M., Winkler, S., Lipinski, K.A., and Tomancak, P. (2009). A toolkit for high-throughput, cross-species gene engineering in Drosophila. Nature Methods 6, 435–437.

Ejsmont, R.K., Ahlfeld, P., Pozniakovsky, A., Stewart, A.F., Tomancak, P., and Sarov, M. (2011). Recombination-mediated genetic engineering of large genomic DNA transgenes. Methods Mol. Biol. 772, 445–458.

Ephrussi, A., Dickinson, L.K., and Lehmann, R. (1991). Oskar organizes the germ plasm and directs localization of the posterior determinant nanos. Cell 66, 37–50.

Fernandes, J., Bate, M., and VijayRaghavan, K. (1991). Development of the indirect flight muscles of Drosophila. Development 113, 67–77.

Fernandes, J.J., Atreya, K.B., Desai, K.M., Hall, R.E., Patel, MD., Desai, A.A., Benham, A.E., Mable, J.L., and Straessle, J.L. (2005). A dominant negative form of Rac1 affects myogenesis of adult thoracic muscles in Drosophila. Developmental Biology 285, 11–27.

Fessler, L.I., Campbell, A.G., Duncan, K.G., and Fessler, J.H. (1987). Drosophila laminin: characterization and localization. Journal of Cell Biology 105, 2383–2391.

Fischer, M., Haase, I., Simmeth, E., Gerisch, G., and Müller-Taubenberger, A. (2004). A brilliant monomeric red fluorescent protein to visualize cytoskeleton dynamics in Dictyostelium. FEBS Letters 577, 227–232.

Formstecher, E., Aresta, S., Collura, V., Hamburger, A., Meil, A., Trehin, A., Reverdy, C., Betin, V., Maire, S., Brun, C., et al. (2005). Protein interaction mapping: a Drosophila case study. Genome Research 15, 376–384.

Forrest, K.M., and Gavis, E.R. (2003). Live imaging of endogenous RNA reveals a diffusion and entrapment mechanism for nanos mRNA localization in Drosophila. Current Biology 13, 1159–1168.

Garrison, E., and Marth, G. (2012). Haplotype-based variant detection from short-read sequencing. 1–9.

Gavin, A.-C., Bösche, M., Krause, R., Grandi, P., Marzioch, M., Bauer, A., Schultz, J., Rick, J.M., Michon, A.-M., Cruciat, C.-M., et al. (2002). Functional organization of the yeast proteome by systematic analysis of protein complexes. Nature 415, 141–147.

Giot, L., Bader, J.S., Brouwer, C., Chaudhuri, A., Kuang, B., Li, Y., Hao, Y.L., Ooi, C.E., Godwin, B., Vitols, E., et al. (2003). A protein interaction map of Drosophila melanogaster. Science 302, 1727–1736.

Gratz, S.J., Ukken, F.P., Rubinstein, C.D., Thiede, G., Donohue, L.K., Cummings, A.M., and O’Connor-Giles, K.M. (2014). Highly specific and efficient CRISPR/Cas9-catalyzed homology-directed repair in Drosophila. Genetics 196, 961–971.

Graveley, B., Brooks, A., Carlson, J., and Duff, M. (2010). The developmental transcriptome of Drosophila melanogaster. Nature.

Guruharsha, K.G., Rual, J.-F., Zhai, B., Mintseris, J., Vaidya, P., Vaidya, N., Beekman, C., Wong, C., Rhee, D.Y., Cenaj, O., et al. (2011). A Protein Complex Network of Drosophila melanogaster. Cell 147, 690–703.

Gutjahr, T., Patel, N.H., Li, X., Goodman, C.S., and Noll, M. (1993). Analysis of the gooseberry locus in Drosophila embryos: gooseberry determines the cuticular pattern and activates gooseberry neuro. Development 118, 21–31.

Haecker, A., Bergman, M., Neupert, C., Moussian, B., Luschnig, S., Aebi, M., and Mannervik, M. (2008). Wollknauel is required for embryo patterning and encodes the Drosophila ALG5 UDP-glucose:dolichyl-phosphate glucosyltransferase. Development 135, 1745–1749.

Hammonds, A.S., Bristow, C.A., Fisher, W.W., Weiszmann, R., Wu, S., Hartenstein, V., Kellis, M., Yu, B., Frise, E., and Celniker, S.E. (2013). Spatial expression of transcription factors in Drosophila embryonic organ development. Genome Biology 14, R140.

Hartenstein, V., Younossi-Hartenstein, A., Lovick, J.K., Kong, A., Omoto, J.J., Ngo, K.T., and Viktorin, G. (2015). Lineage-associated tracts defining the anatomy of the Drosophila first instar larval brain. Developmental Biology 1–26.

He, H., and Noll, M. (2013). Differential and redundant functions of gooseberry and gooseberry neuro in the central nervous system and segmentation of the Drosophila embryo. Developmental Biology 382, 209–223.

Hein, M., Hubner, N., Poser, I., Cox, J., Nagaraj, N., Toyoda, Y., Gak, I., Weisswange, I., Mansfeld, J., Buchholz, F., et al. (2015). A human interactome in three quantitative dimensions organized by stoichiometries and abundances. Cell *in press*.

Ho, Y., Gruhler, A., Heilbut, A., Bader, G.D., and Moore, L. (2002). Systematic identification of protein complexes in Saccharomyces cerevisiae by mass spectrometry. Nature 415, 180–183.

Hubner, N.C., Bird, A.W., Cox, J., Splettstoesser, B., Bandilla, P., Poser, I., Hyman, A., and Mann, M. (2010). Quantitative proteomics combined with BAC TransgeneOmics reveals in vivo protein interactions. The Journal of Cell Biology 189, 739–754.

Huisken, J., and Stainier, D.Y.R. (2007). Even fluorescence excitation by multidirectional selective plane illumination microscopy (mSPIM). Opt Lett 32, 2608–2610.

Huisken, J., Swoger, J., Del Bene, F., Wittbrodt, J., and Stelzer, E.H.K. (2004). Optical sectioning deep inside live embryos by selective plane illumination microscopy. Science 305, 1007–1009.

Izumi, Y., Yanagihashi, Y., and Furuse, M. (2012). A novel protein complex, Mesh-Ssk, is required for septate junction formation in the Drosophila midgut. Journal of Cell Science 125, 4923–4933.

Jambor, H., Brunel, C., and Ephrussi, A. (2011). Dimerization of oskar 3’ UTRs promotes hitchhiking for RNA localization in the Drosophila oocyte. Rna 17, 2049–2057.

Jambor, H., Mueller, S., Bullock, S.L., and Ephrussi, A. (2014). A stem-loop structure directs oskar mRNA to microtubule minus ends. Rna 20, 429–439.

Jambor, H., Surendranath, V., Kalinka, A.T., Mejstrik, P., Saalfeld, S., and Tomancak, P. (2015). Systematic imaging reveals features and changing localization of mRNAs in Drosophila development. eLife 4, e05003.

Jenny, A., Hachet, O., Závorszky, P., Cyrklaff, A., Weston, M.D.J., Johnston, D.S., Erdélyi, M., and Ephrussi, A. (2006). A translation-independent role of oskar RNA in early Drosophila oogenesis. Development 133, 2827–2833.

Kadrmas, J.L., Smith, M.A., Clark, K.A., Pronovost, S.M., Muster, N., Yates, J.R., and Beckerle, M.C. (2004). The integrin effector PINCH regulates JNK activity and epithelial migration in concert with Ras suppressor 1. Journal of Cell Biology 167, 1019–1024.

Katzemich, A., Kreisköther, N., Alexandrovich, A., Elliott, C., Schöck, F., Leonard, K., Sparrow, J., and Bullard, B. (2012). The function of the M-line protein obscurin in controlling the symmetry of the sarcomere in the flight muscle of Drosophila. Journal of Cell Science 125, 3367–3379.

Keilhauer, E.C., Hein, M.Y., and Mann, M. (2014). Accurate Protein Complex Retrieval by Affinity Enrichment Mass Spectrometry (AE-MS) Rather than Affinity Purification Mass Spectrometry (AP-MS). Molecular & Cellular Proteomics 14, 120–135.

Kim-Ha, J., Kerr, K., and Macdonald, P.M. (1995). Translational regulation of oskar mRNA by bruno, an ovarian RNA-binding protein, is essential. Cell 81, 403–412.

Kim-Ha, J., Smith, J.L., and Macdonald, P.M. (1991). oskar mRNA is localized to the posterior pole of the Drosophila oocyte. Cell 66, 23–35.

Krogan, N.J., Cagney, G., Yu, H., Zhong, G., Guo, X., Ignatchenko, A., Li, J., Pu, S., Datta, N., Tikuisis, A.P., et al. (2006). Global landscape of protein complexes in the yeast Saccharomyces cerevisiae. Nature 440, 637–643.

Kvon, E.Z., Kazmar, T., Stampfel, G., Yáñez-Cuna, J.O., Pagani, M., Schernhuber, K., Dickson, B.J., and Stark, A. (2014). Genome-scale functional characterization of Drosophila developmental enhancers in vivo. Nature 512, 91–95.

Langmead, B., and Salzberg, S.L. (2012). Fast gapped-read alignment with Bowtie 2. Nature Methods 9, 357–359.

Leiss, D., Hinz, U., Gasch, A., Mertz, R., and Renkawitz-Pohl, R. (1988). Beta 3 tubulin expression characterizes the differentiating mesodermal germ layer during Drosophila embryogenesis. Development 104, 525–531.

Lécuyer, E., Yoshida, H., Parthasarathy, N., Alm, C., Babak, T., Cerovina, T., Hughes, T.R., Tomancak, P., and Krause, H.M. (2007). Global analysis of mRNA localization reveals a prominent role in organizing cellular architecture and function. Cell 131, 174–187.

Li, H., Handsaker, B., Wysoker, A., Fennell, T., Ruan, J., Homer, N., Marth, G., Abecasis, G., Durbin, R., 1000 Genome Project Data Processing Subgroup (2009). The Sequence Alignment/Map format and SAMtools. Bioinformatics 25, 2078–2079.

Lovick, J.K., Ngo, K.T., Omoto, J.J., Wong, D.C., Nguyen, J.D., and Hartenstein, V. (2013). Postembryonic lineages of the Drosophila brain_ I. Development of the lineage-associated fiber tracts. Developmental Biology 384, 228–257.

Lowe, N., Rees, J.S., Roote, J., Ryder, E., Armean, I.M., Johnson, G., Drummond, E., Spriggs, H., Drummond, J., Magbanua, J.P., et al. (2014). Analysis of the expression patterns, subcellular localisations and interaction partners of Drosophila proteins using a pigP protein trap library. Development 141, 3994–4005.

Meacock, P.A., and Cohen, S.N. (1980). Partitioning of bacterial plasmids during cell division: a cis-acting locus that accomplishes stable plasmid inheritance. Cell 20, 529–542.

Metcalf, W.W., Jiang, W., Daniels, L.L., Kim, S.K., Haldimann, A., and Wanner, B.L. (1996). Conditionally replicative and conjugative plasmids carrying lacZ alpha for cloning, mutagenesis, and allele replacement in bacteria. Plasmid 35, 1–13.

Morin, X., Daneman, R., Zavortink, M., and Chia, W. (2001). A protein trap strategy to detect GFP-tagged proteins expressed from their endogenous loci in Drosophila. Proceedings of the National Academy of Sciences of the United States of America 98, 15050–15055.

Nagarkar-Jaiswal, S., Lee, P.-T., Campbell, M.E., Chen, K., Anguiano-Zarate, S., Gutierrez, M.C., Busby, T., Lin, W.-W., He, Y., Schulze, K.L., et al. (2015). A library of MiMICs allows tagging of genes and reversible, spatial and temporal knockdown of proteins in Drosophila. eLife 4.

Neuman-Silberberg, F.S., and Schüpbach, T. (1993). The Drosophila dorsoventral patterning gene gurken produces a dorsally localized RNA and encodes a TGF alphalike protein. Cell 75, 165–174.

Ni, J.-Q., Zhou, R., Czech, B., Liu, L.-P., Holderbaum, L., Yang-Zhou, D., Shim, H.-S., Tao, R., Handler, D., Karpowicz, P., et al. (2011). A genome-scale shRNA resource for transgenic RNAi in Drosophila. Nature Methods 8, 405–407.

Nongthomba, U., Pasalodos-Sanchez, S., Clark, S., Clayton, J., and Sparrow, J. (2001). Expression and function of the Drosophila ACT88F actin isoform is not restricted to the indirect flight muscles. J Muscle Res Cell Motil 22, 111–119.

Oas, S.T., Bryantsev, A.L., and Cripps, R.M. (2014). Arrest is a regulator of fiber-specific alternative splicing in the indirect flight muscles of Drosophila. The Journal of Cell Biology 206, 895–908.

Pereanu, W. (2006). Neural Lineages of the Drosophila Brain: A Three-Dimensional Digital Atlas of the Pattern of Lineage Location and Projection at the Late Larval Stage. Journal of Neuroscience 26, 5534–5553.

Pédelacq, J.-D., Cabantous, S., Tran, T., Terwilliger, T.C., and Waldo, G.S. (2005). Engineering and characterization of a superfolder green fluorescent protein. Nature Biotechnology 24, 79–88.

Pietzsch, T., Saalfeld, S., Preibisch, S., and Tomancak, P. (2015). BigDataViewer: visualization and processing for large image data sets. Nature Methods 12, 481–483.

Port, F., Chen, H.-M., Lee, T., and Bullock, S.L. (2014). Optimized CRISPR/Cas tools for efficient germline and somatic genome engineering in Drosophila. Proceedings of the National Academy of Sciences 111, E2967–E2976.

Preibisch, S., Amat, F., Stamataki, E., Sarov, M., Singer, R.H., Myers, E., and Tomancak, P. (2014). Efficient Bayesian-based multiview deconvolution. Nature Methods 11, 645–648.

Preibisch, S., Saalfeld, S., and Tomancak, P. (2009). Globally optimal stitching of tiled 3D microscopic image acquisitions. Bioinformatics 25, 1463–1465.

Quiñones-Coello, A.T., Petrella, L.N., Ayers, K., Melillo, A., Mazzalupo, S., Hudson, A.M., Wang, S., Castiblanco, C., Buszczak, M., Hoskins, R.A., et al. (2007). Exploring strategies for protein trapping in Drosophila. Genetics 175, 1089–1104.

Razzaq, A., Robinson, I., McMahon, H., Skepper, J., Su, Y., Zelhof, A., Jackson, A., Gay, N., and O’Kane, C. (2001). Amphiphysin is necessary for organization of the excitation-contraction coupling machinery of muscles, but not for synaptic vesicle endocytosis in Drosophila. Genes & Development 15, 2967.

Rørth, P. (1998). Gal4 in the Drosophila female germline. Mechanisms of Development 78, 113–118.

Rørth, P., Szabo, K., Bailey, A., Laverty, T., Rehm, J., Rubin, G.M., Weigmann, K., Milán, M., Benes, V., Ansorge, W., et al. (1998). Systematic gain-of-function genetics in Drosophila. Development 125, 1049–1057.

Sarov, M., Murray, J.I., Schanze, K., Pozniakovski, A., Niu, W., Angermann, K., Hasse, S., Rupprecht, M., Vinis, E., Tinney, M., et al. (2012). A genome-scale resource for in vivo tag-based protein function exploration in C. elegans. Cell 150, 855–866.

Schindelin, J., Arganda-Carreras, I., Frise, E., Kaynig, V., Longair, M., Pietzsch, T., Preibisch, S., Rueden, C., Saalfeld, S., Schmid, B., et al. (2012). Fiji: an open-source platform for biological-image analysis. Nature Methods 9, 676–682.

Schmied, C., Stamataki, E., and Tomancak, P. (2014). Open-source solutions for SPIMage processing. Methods Cell Biol. 123, 505–529.

Schmied, C., Steinbach, P., Pietzsch, T., Preibisch, S., and Tomancak, P. (2015). An automated workflow for parallel processing of large multiview SPIM recordings. arXiv:1507.085 75, 1–13.

Schnorrer, F., Kalchhauser, I., and Dickson, B. (2007). The transmembrane protein Kon-tiki couples to Dgrip to mediate myotube targeting in Drosophila. Developmental Cell 12, 751–766.

Schnorrer, F., Schönbauer, C., Langer, C.C.H., Dietzl, G., Novatchkova, M., Schernhuber, K., Fellner, M., Azaryan, A., Radolf, M., Stark, A., et al. (2010). Systematic genetic analysis of muscle morphogenesis and function in Drosophila. Nature 464, 287–291.

Schönbauer, C., Distler, J., Jährling, N., Radolf, M., Dodt, H.-U., Frasch, M., and Schnorrer, F. (2011). Spalt mediates an evolutionarily conserved switch to fibrillar muscle fate in insects. Nature 479, 406–409.

Sokolovski, M., Bhattacherjee, A., Kessler, N., Levy, Y., and Horovitz, A. (2015). Thermodynamic Protein Destabilization by GFP Tagging: A Case of Interdomain Allostery. Biophysical Journal 109, 1157–1162.

Spletter, M.L., and Schnorrer, F. (2014). Transcriptional regulation and alternative splicing cooperate in muscle fiber-type specification in flies and mammals. Experimental Cell Research 321, 90–98.

Spletter, M.L., Barz, C., Yeroslaviz, A., Schönbauer, C., Ferreira, I.R.S., Sarov, M., Gerlach, D., Stark, A., Habermann, B.H., and Schnorrer, F. (2015). The RNA-binding protein Arrest (Bruno) regulates alternative splicing to enable myofibril maturation in Drosophila flight muscle. EMBO Rep 16, 178–191.

St Johnston, D. (2012). Using mutants, knockdowns, and transgenesis to investigate gene function in Drosophila. WIREs Dev Biol 2, 587–613.

Sun, B., Xu, P., and Salvaterra, P.M. (1999). Dynamic visualization of nervous system in live Drosophila. Proceedings of the National Academy of Sciences of the United States of America 96, 10438–10443.

Tomancak, P., Beaton, A., Weiszmann, R., Kwan, E., Shu, S., Lewis, S.E., Richards, S., Ashburner, M., Hartenstein, V., Celniker, S.E., et al. (2002). Systematic determination of patterns of gene expression during Drosophila embryogenesis. Genome Biology 3, RESEARCH0088.

Tomancak, P., Berman, B.P., Beaton, A., Weiszmann, R., Kwan, E., Hartenstein, V., Celniker, S.E., and Rubin, G.M. (2007). Global analysis of patterns of gene expression during Drosophila embryogenesis. Genome Biology 8, R145.

Tracey, W.D., Wilson, R.I., Laurent, G., and Benzer, S. (2003). painless, a Drosophila gene essential for nociception. Cell 113, 261–273.

Trcek, T., Grosch, M., York, A., Shroff, H., Lionnet, T., and Lehmann, R. (2015). Drosophila germ granules are structured and contain homotypic mRNA clusters. Nature Communications 6, 7962.

Tseng, A.-S.K., and Hariharan, I.K. (2002). An overexpression screen in Drosophila for genes that restrict growth or cell-cycle progression in the developing eye. Genetics 162, 229–243.

Tu, Y., Huang, Y., Zhang, Y., Hua, Y., and Wu, C. (2001). A new focal adhesion protein that interacts with integrin-linked kinase and regulates cell adhesion and spreading. Journal of Cell Biology 153, 585–598.

Urbach, R., and Technau, G.M. (2003). Molecular markers for identified neuroblasts in the developing brain of Drosophila. Development 130, 3621–3637.

Vaskova, M., Bentley, A.M., Marshall, S., Reid, P., Thummel, C.S., and Andres, A.J. (2000). Genetic analysis of the Drosophila 63F early puff. Characterization of mutations in E63-1 and maggie, a putative Tom22. Genetics 156, 229–244.

Venken, K.J.T., and Bellen, H.J. (2012). Genome-Wide Manipulations of Drosophila melanogaster with Transposons, Flp Recombinase, and ΦC31 Integrase. Methods Mol. Biol. 859, 203–228.

Venken, K.J.T., Carlson, J.W., Schulze, K.L., Pan, H., He, Y., Spokony, R., Wan, K.H., Koriabine, M., de Jong, P.J., White, K.P., et al. (2009). Versatile P[acman] BAC libraries for transgenesis studies in Drosophila melanogaster. Nature Methods 6, 431–434.

Venken, K.J.T., He, Y., Hoskins, R.A., and Bellen, H.J. (2006). P[acman]: a BAC transgenic platform for targeted insertion of large DNA fragments in D. melanogaster. Science 314, 1747–1751.

Venken, K.J.T., Schulze, K.L., Haelterman, N.A., Pan, H., He, Y., Evans-Holm, M., Carlson, J.W., Levis, R.W., Spradling, A.C., Hoskins, R.A., et al. (2011). MiMIC: a highly versatile transposon insertion resource for engineering Drosophila melanogaster genes. Nature Methods 8, 737–743.

Vernes, S.C. (2014). Genome wide identification of fruitless targets suggests a role in upregulating genes important for neural circuit formation. Sci Rep 4, 4412.

Vigoreaux, J., Saide, J., Valgeirsdottir, K., and Pardue, M. (1993). Flightin, a novel myofibrillar protein of Drosophila stretch-activated muscles. Journal of Cell Biology 121, 587.

Wangler, M.F., Yamamoto, S., and Bellen, H.J. (2015). Fruit flies in biomedical research. Genetics 199, 639–653.

Weitkunat, M., and Schnorrer, F. (2014). A guide to study Drosophila muscle biology. Methods 68, 2–14.

Weitkunat, M., Kaya-Copur, A., Grill, S.W., and Schnorrer, F. (2014). Tension and force-resistant attachment are essential for myofibrillogenesis in Drosophila flight muscle. Curr Biol 24, 705–716.

Younossi-Hartenstein, A., Salvaterra, P.M., and Hartenstein, V. (2002). Early development of theDrosophila brain: IV. Larval neuropile compartments defined by glial septa. The Journal of Comparative Neurology 455, 435–450.

Zhang, X., Koolhaas, W.H., and Schnorrer, F. (2014). A versatile two-step CRISPR- and RMCE-based strategy for efficient genome engineering in Drosophila. G3 (Bethesda) 4, 2409–2418.

